# Febrile temperature enhances *Plasmodium falciparum* and neutrophil adhesion by disrupting the endothelial glycocalyx

**DOI:** 10.1101/2025.09.07.674757

**Authors:** Viola Introini, Rory KM Long, Olawunmi Rashidat Oyerinde, María Gestal-Mato, Silvia Sanz Sender, Iván de Jesús Rizo Álvarez, Leticia Hartmann, Frank Stein, Gyu Min Hwang, Borja Lopez Gutierrez, Karl B Seydel, Gretchen Birbeck, Maria Bernabeu

## Abstract

Fever, a universal host defense response in infection and inflammation, paradoxically contributes to neurological complications in malaria. Febrile temperatures are known to enhance parasite virulence protein expression, but direct effects on the human endothelium remain unknown. We found that a 1-hour exposure to 40 °C, representative of fever in children with cerebral malaria, increased adhesion of *Plasmodium falciparum*-infected red blood cells and neutrophils to 3D brain microvascular models displaying a wide wall shear stress gradient. Mechanistically, this brief hyperthermia triggered rapid endothelial glycocalyx shedding, exposing endothelial receptors for binding. This response was more pronounced in brain than in pulmonary microvessels, revealing a greater vulnerability of the cerebral vasculature to fever. Pharmacological inhibition of matrix metalloproteinase activity preserved glycocalyx integrity and abolished the temperature-induced increase in adhesion. These findings identify fever as a host-specific amplifier of malaria-associated microvascular pathology, highlighting the importance of antipyretic strategies to mitigate disease severity.

## Main

Fever is an ancient and metabolically costly host response to infection and inflammation, conserved across vertebrates. This tightly regulated process is initiated when injury- and pathogen-associated molecular patterns trigger immune responses and release of prostaglandin^1,2^, a lipid effector molecule that activates thermoregulatory neuronal circuits in the hypothalamus to rapidly elevate core body temperature^3^. Despite these protective functions, fever is not universally beneficial to the host: in diseases such as sepsis^4^, neurological injury^5^, and malaria^6^, fever suppression improves outcomes.

Fever is a hallmark of malaria infection and a consistent predictor of severity^7,8^. The detrimental effects of fever are exemplified by the reduction in seizures following aggressive antipyretic treatment in children with cerebral malaria^9^. Central to severe malaria pathogenesis is the cytoadherence of *Plasmodium falciparum*-infected red blood cells (iRBCs) to the vascular endothelium^10^, mediated by clonally variant *P. falciparum* erythrocyte membrane protein 1 (PfEMP1) proteins^11,12^. Binding of PfEMP1 to endothelial protein C receptor (EPCR) and intercellular adhesion molecule 1 (ICAM-1) is strongly linked with severe disease^13^. On the parasite side, febrile temperatures profoundly reshape iRBC biology^14^ inducing cytoskeletal remodelling^15^, and accelerating PfEMP1 surface expression,^16–18,14^, changes that translate into rapid increase in iRBC stiffness^19^, adhesion^20–22^, and endothelial contact^18^. At the immune level, fever mobilises host systemic innate and adaptive immune responses^23^, promoting vascular recruitment of neutrophils and T cells^24^, as well as cytokine release^25^. Together, these observations suggest that fever acts on both iRBCs and immune cells and amplifies vascular interactions. Yet the endothelial response to febrile temperatures remains poorly understood — a critical gap for host-directed therapeutic strategies. Here, we employ bioengineered human brain and pulmonary 3D microvascular models to dissect how febrile temperatures regulate *Plasmodium falciparum* and neutrophil adhesion.

### Paediatric patients experience high fever despite antipyretic treatment

To inform the febrile conditions for *in vitro* experiments, we analysed continuous temperature recordings from 146 children enrolled in a randomised controlled trial of aggressive antipyretic treatment for malarial infection in Malawi. All participants presented Central Nervous System (CNS) symptoms including impaired consciousness and/or seizures, and 72–83% fulfilled the clinical definition of cerebral malaria (CM)^6,9^. Core temperature was measured every 2-4 minutes over 72 h using a monitoring patch. Patients received either standard-of-care or aggressive antipyretic therapy with prophylactic acetaminophen and ibuprofen as described in Methods. Despite antipyretic therapy, 25% of patients reached body temperatures between 39-40 °C, and 10% reached ≥ 40 °C (**Fig. 1a**). The average fever duration was 1 h for ≥ 40 °C and 1.7 h for ≥ 39 °C, with some episodes persisting for several hours (**Fig. 1b**). These findings demonstrate that short episodes of fever are common in CM patients, even under current WHO-recommended fever management.

**Fig. 1.**
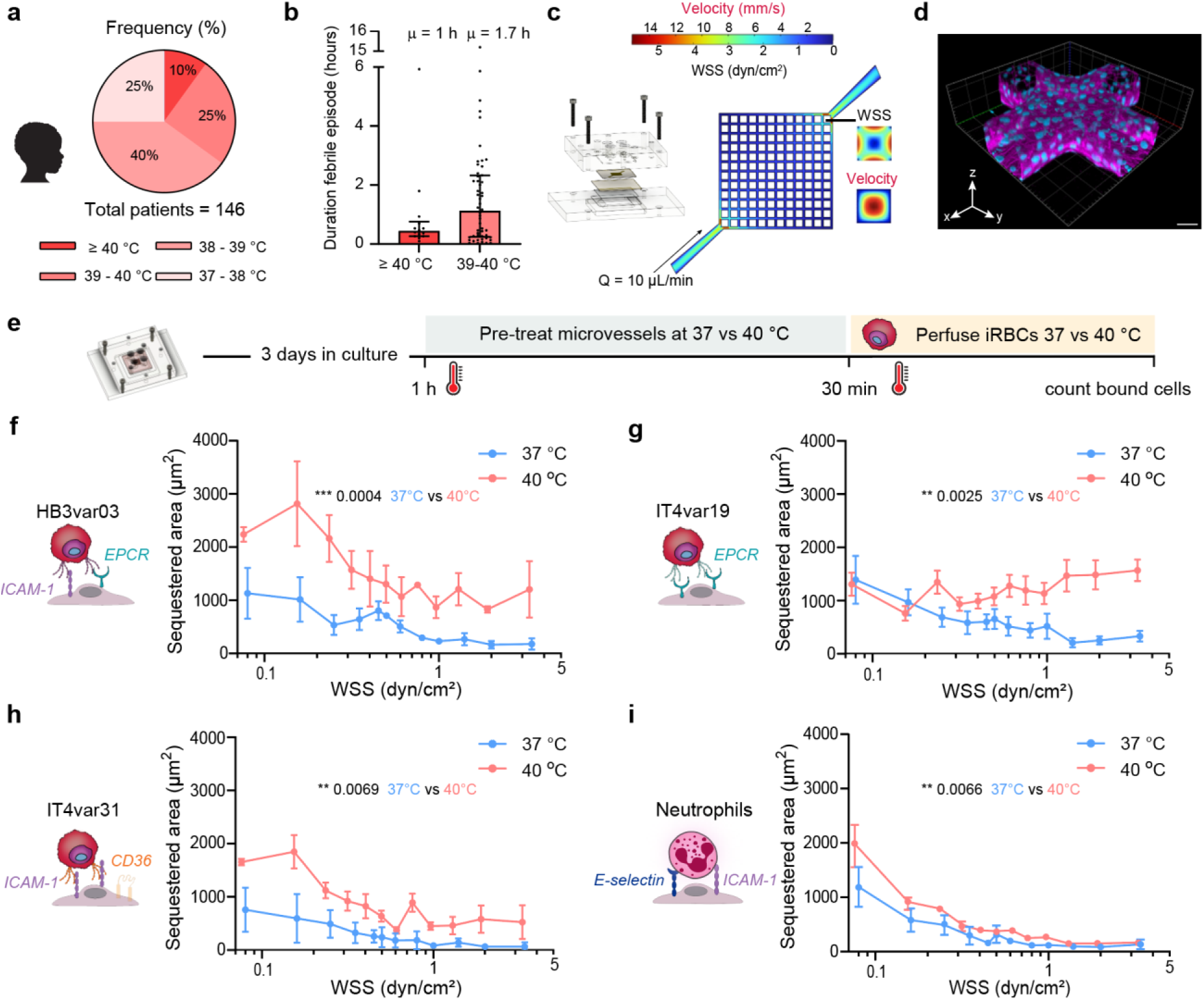
Fever in paediatric cerebral malaria patients and temperature-dependent *P. falciparum* iRBC and neutrophil binding to 3D microvessels. **a,** Distribution of maximum body temperatures recorded during hospital admission in 146 paediatric malaria cases from Malawi treated with antipyretics (see Methods). **b,** Duration of fever episodes with temperatures ≥ 40 °C or between 39 °C and 40 °C. Bars represent median and interquartile range; μ indicates the mean. **c,** Simulation of mid-plane flow velocity in the microvascular grid network before collagen remodelling by HBMECs (see Methods). Inset, lumen cross-sectional flow velocity and wall shear stress (WSS) at the first branch (black line). **d,** Volumetric reconstruction of a microvessel network cross-section immunostained for VE-cadherin (magenta) with nuclei counterstained with DAPI (blue). Scale bar, 50 μm. **e,** Schematic of the binding assay workflow. **f–h,** Schematic of receptor binding and quantification of iRBC sequestration area across WSS range following 1 h incubation of microvessels at 37 °C or 40 °C across the WSS range for **f**, HB3var03, **g**, IT4var19 and **h**, IT4var31. **i,** Quantification of neutrophil adhesion following 1 h incubation at 37 °C or 40 °C across the WSS range. In **f–i**, *n* = 4–7 independent biological replicates. Dots represent the mean and error bars represent the s.e.m.. Statistical comparisons of sequestration areas between 37 °C and 40 °C were performed using a LMM to account for the dependence of binding on WSS.

### Febrile temperature increases iRBC and neutrophil adhesion to 3D brain microvessels

To determine whether hyperthermia influences *P. falciparum* binding to endothelial cells, we employed a bioengineered 3D microvascular model that replicates flow-driven cytoadhesion. The platform consists of a microfluidic network of 13x13 channels of 120 µm diameter embedded in a collagen hydrogel, recapitulating a wide range of flow velocities spanning 0.4–15 mm/s – comparable to those found in the arteriovenous microcirculation (0.4–8 mm/s)^26,27^ **(Fig. 1c**), and equivalent to a wall shear stress (WSS) gradient between 0.08 and 3.4 dyn/cm^2^ (**Extended Data Fig. 1a**). Human brain microvascular endothelial cells (HBMECs) are seeded into the network, and after 3 days in culture, they form fully endothelialised 3D microvessels (**Fig. 1d**). To experimentally model clinical malarial fever episodes, we exposed our engineered 3D human brain microvessel system to 40 °C for 1 h, mimicking the hyperthermic episodes observed in paediatric CM (**Fig. 1b**). Microvessel morphology was largely unaffected by temperature, with no significant changes in endothelial junction organisation, microvessel diameter, endothelial cell area, perimeter, or circularity. Intracellular von Willebrand factor (vWF) mean fluorescence intensity (MFI) showed a slight, non-significant increase, indicating no major heat-induced vessel remodelling in the model (**Extended Data Figs. 2a-f**). Normothermic *P. falciparum*-iRBCs, stained with a fluorescent membrane dye and maintained at 37 °C prior to perfusion, were then introduced into the microvessels for 30 min. Perfusion was performed at either 37 °C (control) or 40 °C, allowing us to distinguish the effects of hyperthermic exposure on the microvessel environment from those of normothermic iRBCs (**Fig. 1e**). Binding was quantified along the outer edges of the microvascular grid, and fluorescent areas with cytoadhered *P. falciparum-*iRBCs were defined as the sequestered area (shown in **Extended Data Figs. 1a,b**). Since temperature influences medium viscosity and density, we first simulated WSS profiles under normothermic (37 °C) and febrile (40 °C) conditions, revealing only minimal differences (**Extended Data Fig. 1c**). Experimental measurements of flow velocity using single-particle tracking and particle image velocimetry (PIV) in the regions of the microvascular network used for binding assays showed no significant temperature-dependent differences (**Extended Data Fig. 1d, e**). Three clonal lines of *P. falciparum-*iRBCs were perfused, expressing predominantly single PfEMP1 variants with well-known binding phenotype to endothelial receptors (**Extended Data Figs. 3a,b**): HB3var03 (dual EPCR and ICAM-1 binder), IT4var19 (EPCR binder), and IT4var31 (dual CD36 and ICAM-1 binder). Statistical significance for all binding experiments in this study was assessed using a linear mixed-effects model (LMM) for the full range of WSS values, accounting for repeated measurements across different regions of accelerating and decelerating flow (see Methods). A WSS threshold sensitivity analysis was performed to test for threshold-dependent behaviour, in which the LMM was refit on cumulative subsets of data, each including all measurements below a progressively increasing WSS cutoff **Extended Data Fig. 4**. All parasite lines showed significantly increased binding at 40 °C. For IT4var19, the WSS threshold sensitivity analysis showed that this effect became more pronounced at higher WSS, reaching significance at the 0.74 dyn/cm^2^ cutoff, which highlights PfEMP1 variant-specific responses to febrile temperature (**Extended Data Fig. 4a**). Temperature did not affect uninfected RBC binding, which remained negligible (**Extended Data Fig. 5a**). As an additional negative control, we used a parasite line expressing var2csa that binds to chondroitin sulfate A, a glycan on syncytiotrophoblasts of the placenta absent on endothelial cells^28^. NF54var2csa showed minimal binding at 37 °C and no increase at 40 °C, supporting that temperature-dependent increase in cytoadhesion requires parasite variants capable of engaging receptors expressed on the endothelial surface (**Extended Data Fig. 5b**). As immune cells engage some of the same endothelial receptors as iRBCs to adhere to the vasculature, including ICAM-1^24^, we next perfused human neutrophils through brain microvessels pre-incubated for 1 h at either 37 °C or 40 °C. Exposure to 40 °C significantly increased neutrophil adhesion across the WSS range (**Fig. 1i**), with very few neutrophils appearing in the collagen surrounding the vessels (**Extended Data Fig. 5c**). Together, these data show that febrile temperature promotes flow-dependent endothelial cytoadhesion of *P. falciparum*-iRBCs and neutrophils.

### EPCR and ICAM-1 mediate temperature-driven iRBC and neutrophil binding without changes in receptor expression

To investigate whether endothelial receptors are responsible for the mechanism underlying increased sequestration at 40 °C, we inhibited *P. falciparum-*iRBC binding using anti-ICAM-1 and anti-EPCR antibodies. Systematic inhibition of ICAM-1, EPCR, or both at 40 °C reduced binding of *P. falciparum* lines HB3var03 andIT4var19 to levels similar or lower to IgG controls at 37 °C (**Figs. 2a,b; Extended Data Figs. 5d,e**). To confirm that the enhanced interaction with endothelial receptors at febrile temperatures was driven by PfEMP1, we incubated iRBCs with the anti-PfEMP1 monoclonal antibody C7, which blocks PfEMP1 interactions with EPCR^29^. C7 antibody treatment significantly inhibited parasite binding at 40 °C (**Fig. 2c**). Altogether demonstrating that PfEMP1-EPCR/ICAM-1 interactions are responsible for iRBC binding under febrile conditions. Next, we treated 3D microvessels with an anti-ICAM-1 blocking antibody before neutrophil perfusion at 40 °C ^24^. ICAM-1 blockade reduced adhesion, with significant inhibition observed when the analysis was restricted to cumulative WSS values up to 0.89 dyn/cm², an effect that was lost when higher-shear measurements were included (**Fig. 2d**, **Extended Data Fig. 4b**). To determine whether febrile temperature increases the expression of ICAM-1 or EPCR and thereby contributes to enhanced iRBC and neutrophil adhesion, we quantified the endothelial proteome using tandem mass tag (TMT)-based quantitative mass spectrometry. Whole-proteome analysis of HBMEC monolayers exposed to 40 °C for 1 h revealed minimal changes in protein expression (**Fig. 2e**). Whereas EPCR and ICAM-1 protein levels remained unchanged after 1 h of febrile temperature exposure (**Extended Data Figs. 6a,b**), only a handful of proteins presented significantly increased levels (e.g., SFSWAP, UQCC1, DPP9) and one showed decreased levels (SNRPD3). These proteins are primarily associated with RNA splicing, mitochondrial function, or inflammasome regulation. Furthermore, the proteomic analysis showed slight increases in proteins related to endothelial dysfunction and heat response, including heparanase (HSPE), heat shock proteins (HSPs), and prostaglandin synthesis pathway enzymes (PTGs), yet not significantly (**Extended Data Figs. 6c-f**). We then further validated these candidates by Western blot to determine if they show further increase over longer periods of febrile temperature exposure. Heparanase protein expression remained unchanged at 1, 6, and 24 h at 40 °C (**Extended Data Fig. 6g**). HSP70 protein abundance increased significantly after 6 and 24 h of exposure to 40 °C but not at earlier timepoints (**Extended Data Fig. 6h**), whereas PTGS2 protein levels were unchanged at early time points and decreased significantly after 24 h (**Extended Data Fig. 6i**). Because the short exposure to febrile temperatures did not cause global changes in protein levels, we next used flow cytometry to quantify potential changes in surface expression of endothelial receptors. No differences in the surface expression of EPCR and ICAM-1 were observed between 37 °C and 40 °C. As HB3var03 can engage both EPCR and ICAM-1, we quantified the proportion of ICAM-1⁺EPCR⁺ double-positive endothelial cells, which also remained unchanged. Only a small percentage of endothelial cells expressed E-selectin, a key mediator of initial neutrophil–endothelial interactions prior to ICAM-1 engagement^30^, and CD36, was undetected, while maintenance of CD31 expression at 40 °C confirmed endothelial cell viability under febrile conditions (**Fig. 2f; Extended Data Figs. 7a–b**). Altogether, these findings indicate that the enhanced *P. falciparum*-iRBC and neutrophil adhesion at 40 °C remains dependent on ICAM-1 and EPCR but cannot be explained by increased endothelial surface levels of these receptors.

**Fig. 2.**
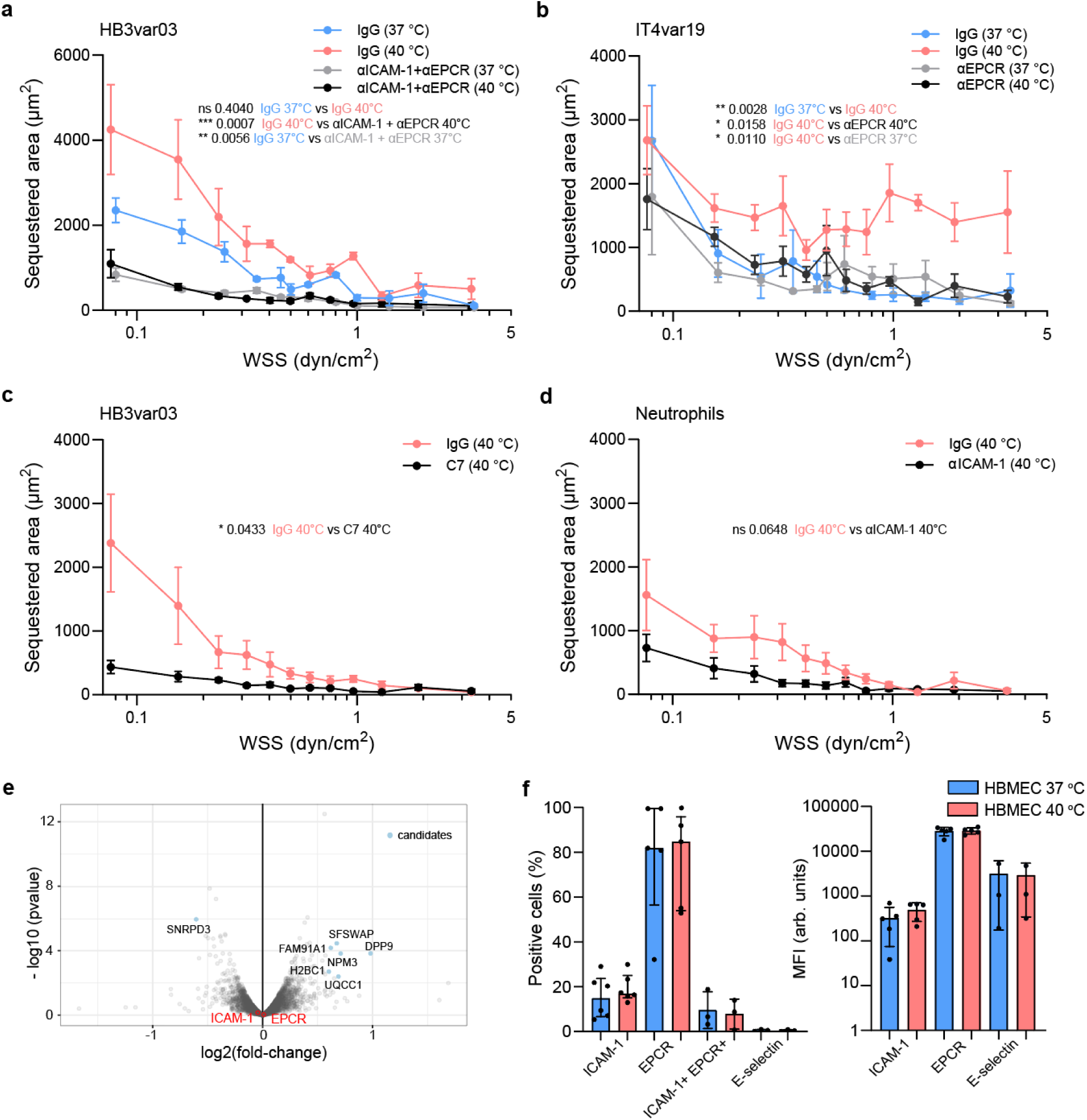
Contribution of endothelial receptors EPCR and ICAM-1 to enhanced cytoadhesion. **a,** HB3var03 iRBC sequestration area across the WSS range following 1 h incubation of microvessels at 37 °C or 40 °C in the presence of IgG isotype control or both anti-EPCR mAb 252 and anti-ICAM-1 mAb 15.2 antibodies. **b,** IT4var19 iRBC sequestration area across the WSS range following 1 h incubation of microvessels at 37 °C or 40 °C in the presence of IgG isotype control or anti-EPCR mAb 252. **c,** HB3var03 iRBC sequestration area across the WSS range in 1-h 40 °C incubated microvessels following 30-minute incubation of iRBCs with IgG isotype control or the anti-CIDRα1 monoclonal antibody C7 against PfEMP1. **d,** Neutrophil adhesion area across the WSS range following 1 h incubation of microvessels at 40 °C in the presence of IgG isotype control or anti-ICAM-1 mAb 15.2. For **a-d**, *n* = 4–6 independent biological replicates. Dots represent the mean and error bars represent the s.e.m. Statistical analysis between sequestration areas was performed using LMM to account for WSS dependence. **e,** Volcano plot of differential protein expression in HBMEC monolayers following 1 h exposure to 37 °C or 40 °C, with significantly regulated proteins labelled. *n* = 3 independent biological replicates. **f,** Percentage of positive cells and mean fluorescence intensity (MFI) of HBMEC surface ICAM-1, EPCR and E-selectin expression, together with the percentage of ICAM-1/EPCR double-positive cells, measured by flow cytometry following 1 h exposure to 37 °C or 40 °C. *n* = 3–6 independent biological replicates. Bars represent mean ± s.d. Statistical analysis was performed using two-sided unpaired Welch’s *t*-tests or Mann–Whitney *U* tests.

### Febrile temperature promotes brain endothelial glycocalyx shedding

The endothelial glycocalyx is a carbohydrate-rich meshwork that covers the luminal surface of the vasculature, limiting endothelial receptor accessibility^31,32^. To determine whether febrile temperature disrupts it, we exposed brain microvessels to 37, 40 °C, or alternatively a control 30-minute neuraminidase (NA) treatment at 37 °C, known to cleave terminal sialic acid residues from membrane glycoproteins^33^. All glycocalyx analyses were performed after 3 days of gravity-driven media on the same day that binding experiments were conducted, as extending microvessel maturation to 7 days did not significantly improve glycocalyx formation, including general sialic acids, stained by wheat germ agglutinin (WGA), or heparan sulfate expression (**Extended Data Figs. 8a,b**). Glycocalyx measurements were acquired from both low and high-shear regions of the microvascular network, and no WSS-dependent significant differences in sialic acid and heparan sulfate were found (**Extended Data Figs. 8c,d**). Febrile temperature induced a reduction in key glycocalyx components including sialic acids and N-acetylglucosamine (WGA; **Fig. 3a**), heparan sulfate (**Fig. 3b**), syndecan-4 (**Fig. 3c**), and α2,6-linked sialic acids detected by the lectin *Sambucus nigra* (**Fig. 3d**). The reduction was, comparable to or even greater than that induced by NA treatment. We further showed by ELISA that Syndecan-4, the most abundant proteoglycan, presented significant shedding, similar to previous observations in experimental CM^34^ (**Fig. 3e**). Because malaria is characterised by recurrent febrile episodes, we next investigated whether a repeated fever peak exacerbates endothelial glycocalyx damage. Among the 17 patients in the CM cohort who experienced two consecutive fever peaks ≥ 40 °C, the mean interval between peaks was 7.8 ± 2.3 h (mean ± s.d.) (**Fig. 3f**). We recreated this fever cycle *in vitro*, with exposure of 3D microvessels to 40 °C for 1 h, followed by a return to 37 °C for 6 h, and a second 1 h peak at 40 °C. Quantification of heparan sulfate staining revealed no additional glycocalyx loss upon the second febrile episode compared with a single febrile episode (**Fig. 3g**). Heparan sulfate did not recover when devices were kept at 37 °C throughout the second cycle, suggesting a lack of glycocalyx recovery within the 7 h time period. Similarly, syndecan-4 shedding remained decreased following the second febrile episode, further arguing against additional glycocalyx degradation on consecutive cycles (**Fig. 3h**).

**Fig. 3.**
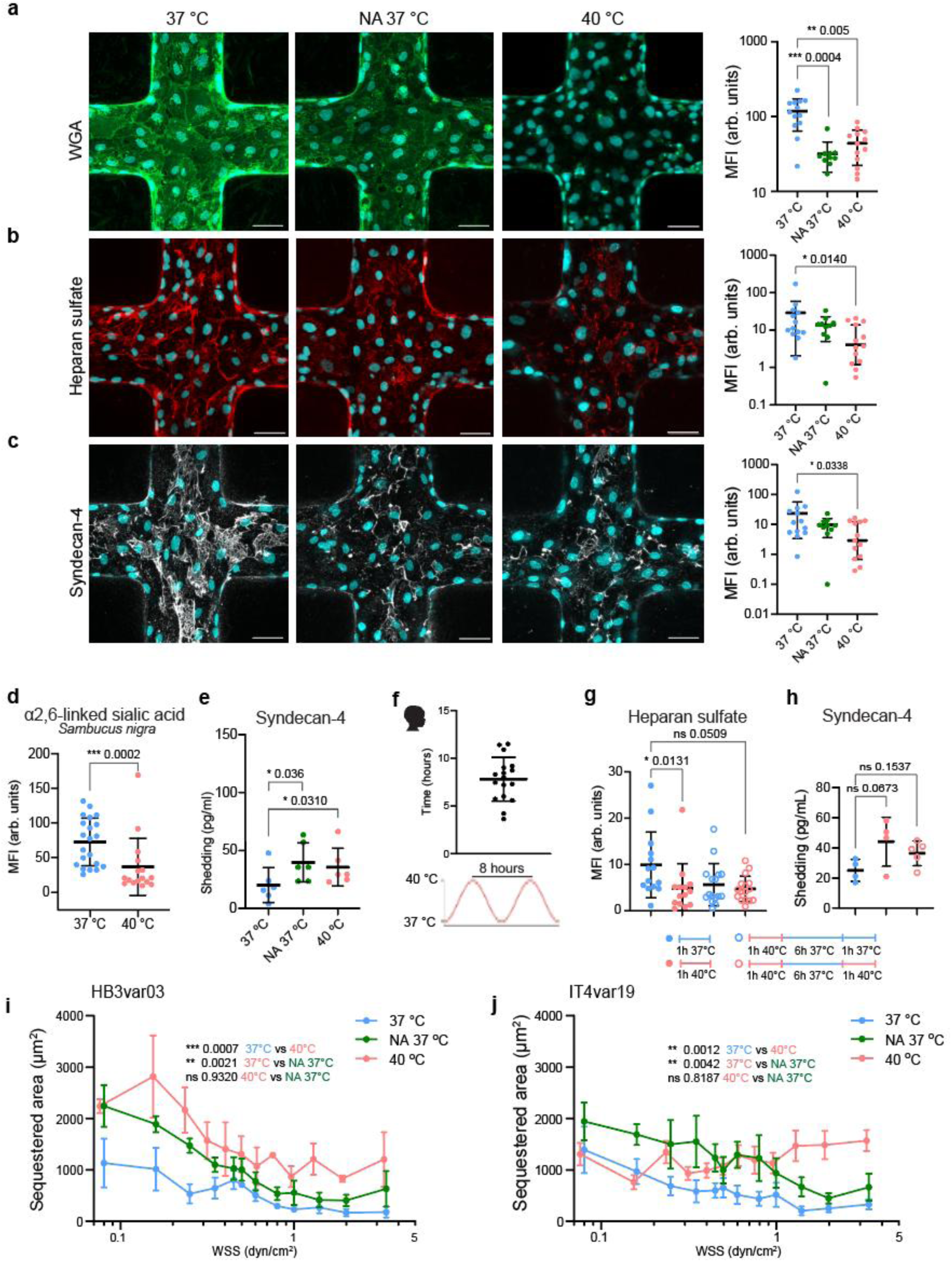
Endothelial glycocalyx shedding in brain microvessels following acute or cycling febrile temperature exposure. **a–c,** Representative immunofluorescence z-stacks of microvessel cross-sections (left) and quantification of mean fluorescence intensity (MFI) normalised to cell number (right) for **a,** sialic acids labelled with FITC-conjugated wheat germ agglutinin (WGA), **b,** heparan sulfate, and **c,** syndecan-4. *n* = 4 microvessels per condition, with each dot representing an ROI. Dot plots show mean ± s.d. Statistical analysis was performed using the Kruskal–Wallis with Dunn’s multiple-comparisons tests. Scale bars, 50 µm. Conditions in **a–c** include microvessels incubated at 37 °C or 40 °C for 1 h, or treated with neuraminidase (NA; 1 U/ml) for 30 min at 37 °C. **d,** MFI of α2,6-linked sialic acid stained with *Sambucus nigra* lectin, normalised to cell number, following 1 h incubation at 37 °C or 40 °C. *n* = 4 microvessels per condition, with each dot representing an ROI (mean ± s.d.). Statistical analysis was done using Mann–Whitney *U* test. **e,** ELISA quantification of shed syndecan-4 in supernatants collected from 3D microvessels following a 1 h incubation at 37 °C or 40 °C, or treated with neuraminidase (NA; 1 U/ml) for 30 min at 37 °C. *n* = 6 microvessels per condition, graph shows mean ± s.d., with each dot representing an independent microvessel batch. Statistical analysis was done using one-way ANOVA with Tukey’s multiple-comparisons test. **f,** Top: average interval between fever episodes reaching ≥ 40 °C. *n* = 17 patients. Dot plot shows mean ± s.d.. Bottom: schematic illustrating the ∼8 h interval between recurrent febrile peaks observed in the paediatric cohort. **g,** MFI of heparan sulfate, normalised to cell number, following treatment with 37 °C for 1 h, 40 °C for 1 h, cycling fever (1 h at 40 °C, followed by 6 h at 37 °C, then a second 1 h incubation at 40 °C), or cycling control (1 h at 40 °C, followed by 6 h at 37 °C, then 1 h at 37 °C). *n* = 4 microvessels per condition, with each dot representing an ROI (mean ± s.d.). Statistical analysis was performed using the Kruskal–Wallis with Dunn’s multiple-comparisons tests. **h,** Syndecan-4 shedding quantified by ELISA in supernatants from 3D brain microvessels following the same temperature conditions in **g**. *n* = 4-5 microvessels per condition, graph shows mean ± s.d., with each dot representing an independent microvessel batch. Statistical analysis was done using one-way ANOVA with Tukey’s multiple-comparisons test. **i,j,** Binding of **i**, HB3var03 and **j**, IT4var19 to 3D brain microvessels treated with NA at 37 °C (*n* = 5) compared with microvessels incubated at 37 °C or 40 °C for 1 h (*n* = 6). Mean is shown as dots and error bars represent the s.e.m. Statistical comparisons of sequestration areas were performed using LMM to account for WSS dependence.

To determine whether glycocalyx degradation causes enhanced *P. falciparum-*iRBC binding, we pretreated 3D microvessels with NA for 30 minutes at 37 °C, followed by perfusion with HB3var03 and IT4var19 lines. Binding was significantly increased for both parasite lines in a LMM, reaching levels comparable to those observed at 40 °C (**Figs. 3i,j**). Incubation with anti-EPCR and ICAM-1 inhibitory antibodies with and without NA treatment, showed a similar reduction of iRBC binding levels, confirming that glycocalyx disruption enhances endothelial receptor engagement by PfEMP1. (**Extended Data Fig. 8e**). Our findings indicate that febrile temperatures induce shedding of endothelial sialic acids and other glycan components that remained unchanged between consecutive fever episodes, thereby facilitating PfEMP1-mediated adhesion.

### Febrile temperature promotes pulmonary endothelial glycocalyx shedding

The pulmonary microvasculature represents another site of pathogenic iRBC cytoadhesion, associated with acute respiratory distress syndrome during severe malarial infection^34,35^. To determine whether temperature-induced glycocalyx shedding is specific to the brain endothelium or also occurs in other vascular beds, we generated a 3D pulmonary microvessel model from primary human pulmonary artery endothelial cells (HPAECs) (**Extended Data Fig. 9a)**^36,37^. Sialic acid and N-acetylglucosamine levels, assessed by WGA staining, were significantly lower in pulmonary microvessels than in brain microvessels at 37°C (**Fig. 4a**); heparan sulfate levels showed a trend for reduced presence but did not reach significance (**Fig. 4b**). Despite these lower baseline levels, pulmonary microvessels exhibited a reduction in sialic acid and N-acetylglucosamine following 1 h incubation at 40 °C, whereas heparan sulfate levels remained unchanged compared with 37 °C, both conditions mirror the effects of NA treatment in 3D pulmonary microvessels (**Figs. 4c,d**). Febrile temperature also increased shedding of syndecan-1 (**Fig. 4e**), a marker of pulmonary vascular pathology in malaria^38,39^. Similar to HBMECs, HPAECs showed no significant differences between 37 °C and 40 °C in the percentage of EPCR^+^, ICAM-1^+^, ICAM-1⁺EPCR⁺, CD31^+^, or CD36^+^ cells, or surface expression MFI of these markers (**Extended Data Figs. 9b,c**). Compared with HBMECs, HPAECs tended to exhibit a lower proportion of EPCR^+^ and CD31^+^ cells and higher surface expression of ICAM-1, although none of these differences reached statistical significance (**Extended Data Figs. 9d,e**). In line with this, both parasite lines showed lower cytoadhesion rates in pulmonary microvessels than in 3D brain microvessels at 37 °C, probably as a consequence of lower pulmonary EPCR expression (**Fig. 4f,g**). Finally, we assessed cytoadhesion of HB3var03- and IT4var19-expressing parasites in pulmonary microvessels incubated at febrile temperatures. Although binding increased at 40 °C for both cell lines, the effect did not reach statistical significance overall. A significant increase was observed for IT4var19 at WSS <0.85 dyn/cm^2^, with this threshold identified using a LMM with WSS threshold sensitivity analysis (**Extended Data Fig. 4c,d**). Therefore, febrile temperature induces glycocalyx shedding in pulmonary microvessels, but to a lesser extent than in brain microvessels, which results in modest effects on parasite cytoadhesion.

**Fig 4.**
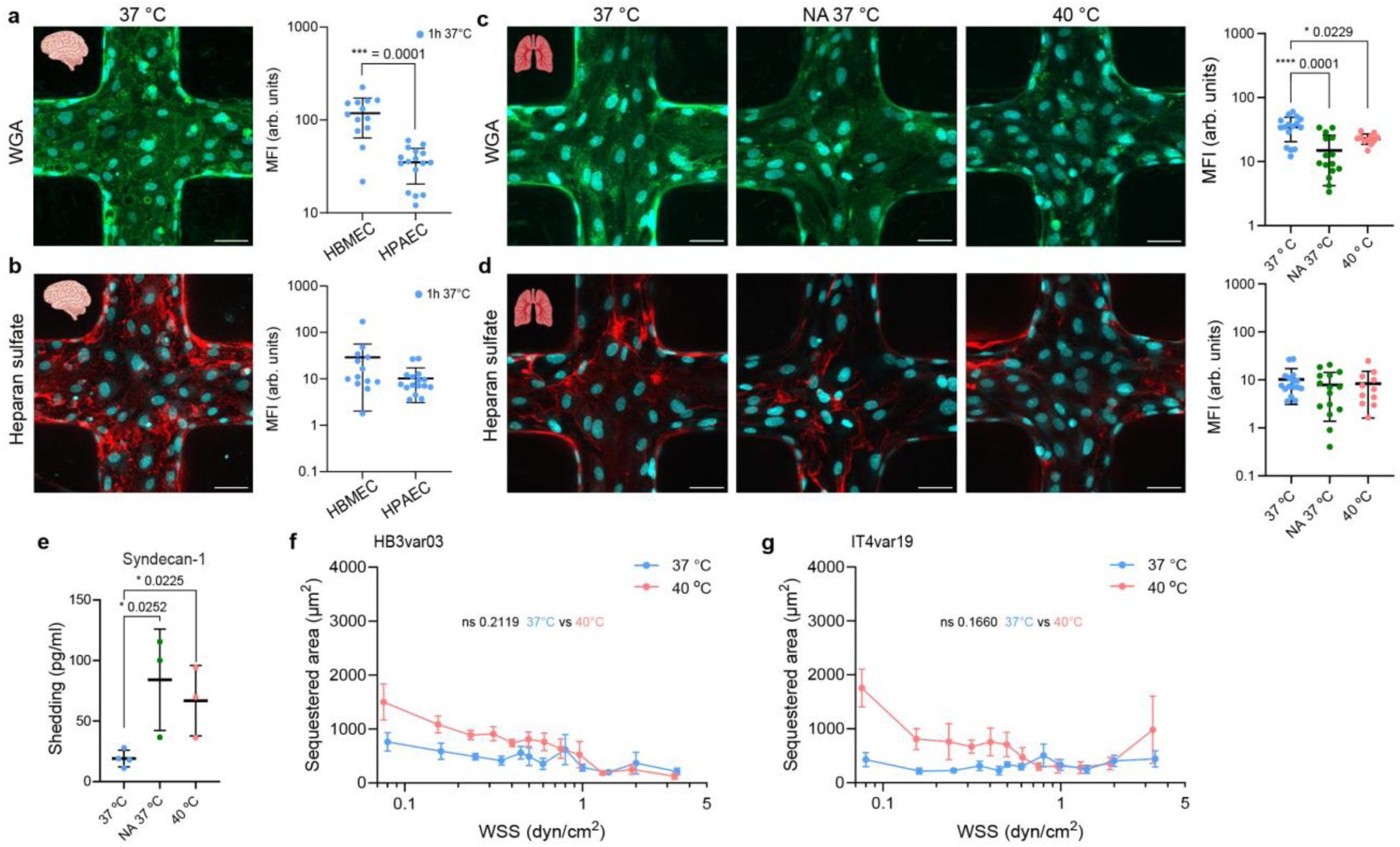
Endothelial glycocalyx shedding in pulmonary microvessels upon 1h-temperature exposure. **a,b,** Representative Immunofluorescence z-stacks of 3D brain microvessel cross-sections (left) and corresponding mean fluorescence intensity (MFI), comparing 3D brain and lung pulmonary microvessels (right). MFI values were normalised to cell number, for sialic acids labelled with **a**, FITC-conjugated wheat germ agglutinin (WGA) and **b**, heparan sulfate between 3D brain and pulmonary microvessels a. *n* = 4 microvessels per condition, with each dot representing an ROI (mean ± s.d.). Statistical analysis was performed using **a**, two-sided unpaired Welch’s *t*-tests and **b**, Mann–Whitney *U* test. Scale bars, 50 µm. **c,d,** Representative immunofluorescence z-stacks of 3D lung microvessel cross-sections (left) and corresponding quantified MFI, normalised to cell number (right), for **c**, WGA and **d**, heparan sulfate in 3D pulmonary microvessels. *n* = 4 microvessels per condition, with each dot representing an ROI (mean ± s.d.). Statistical analysis was performed using the Kruskal–Wallis with Dunn’s multiple-comparisons tests. Scale bars, 50 µm **e,** Syndecan-1 shedding quantified by ELISA in supernatants collected from 3D pulmonary microvessels. *n* = 3-4 microvessels per condition, graph shows mean ± s.d., with each dot representing an independent microvessel batch. Statistical analysis was performed using the Kruskal–Wallis with Dunn’s multiple-comparisons tests. For **c, d, e**, microvessels were incubated at 37 °C or 40 °C for 1 h, or treated with neuraminidase (NA; 1 U/ml) for 30 min at 37 °C. **f,g,** iRBC sequestered areas in 3D pulmonary microvessels following 1-hour 37 °C and 40 °C incubation across WSS for: **f**, HB3var03; **g**, IT4var19. Mean is represented by dots and s.e.m. by error bars. (*n* = 5 independent biological replicates). Statistical analysis between sequestered areas at 37 °C and 40 °C was determined using LMM to account for WSS-dependence.

### Matrix metalloproteinase inhibition preserves the glycocalyx and prevents temperature-induced vascular adhesion

To investigate the mechanisms driving glycocalyx degradation at febrile temperature, we analysed the secretion of angiogenic mediators, inflammatory cytokines and proteolytic enzymes from HBMEC monolayers using a semi-quantitative protein array. Following 1 h incubation at 40 °C, MMP-1 secretion showed the largest and most consistent increase, angiopoietin-2 (ANGPT2) and the endogenous matrix metalloproteinase inhibitor TIMP-2 to a lesser extent but none of these changes reached statistical significance (**Fig. 5a; Extended Data Figs. 10a–c**). No changes were detected in the endothelial-secreted pro-inflammatory cytokines IL-6 and IL-8 or the chemokine MCP-1. As MMP-1 has previously been implicated in endothelial glycocalyx degradation^40^, we next investigated whether febrile temperature increases secreted MMP activity. Measurements of MMP activity in supernatants collected from HBMECs incubated for 1 h at either 37 °C or 40 °C were conducted using a broad MMP fluorogenic peptide substrate. Endothelial cells exposed to 40 °C showed increased MMP activity compared with 37 °C (p = 0.0687) (**Fig. 5b**). Another alternative candidate for glycocalyx shedding is heparanase, a well-established mediator of glycocalyx degradation through cleavage of heparan sulfate in metastatic cancer, viral infection, and sepsis^41–43^. Heparanase levels in HBMEC supernatants were unchanged following 1 h incubation at 37 °C or 40 °C (**Extended Data Fig. 10d**), consistent with our western blot analysis in **Extended Data Fig. 6g**. Notably, broad-spectrum MMP inhibition with batimastat abolished the temperature-dependent increase in MMP activity, with minimal activity levels both at 37 °C and 40 °C (**Fig. 5b**) and prevented the temperature-induced loss of sialic acid, heparan sulfate, and syndecan-4 (**Figs. 5c–e**). Consistent with glycocalyx preservation, batimastat significantly reduced adhesion of both HB3var03- and IT4var19-iRBCs (**Figs. 5f,g; Extended Data Figs. 10e,f**), and neutrophils (**Fig. 5h**), to normothermic levels. Collectively, these findings support the mechanism illustrated in **Fig. 5i**: brief exposure of the vascular endothelium to febrile temperatures promotes MMP-dependent glycocalyx degradation, enhancing vascular adhesion across diverse vascular beds. Pharmacological MMP inhibition preserves glycocalyx integrity and prevents fever-induced adhesion, highlighting this process as a potential host-directed therapeutic target in severe malaria.

**Fig. 5.**
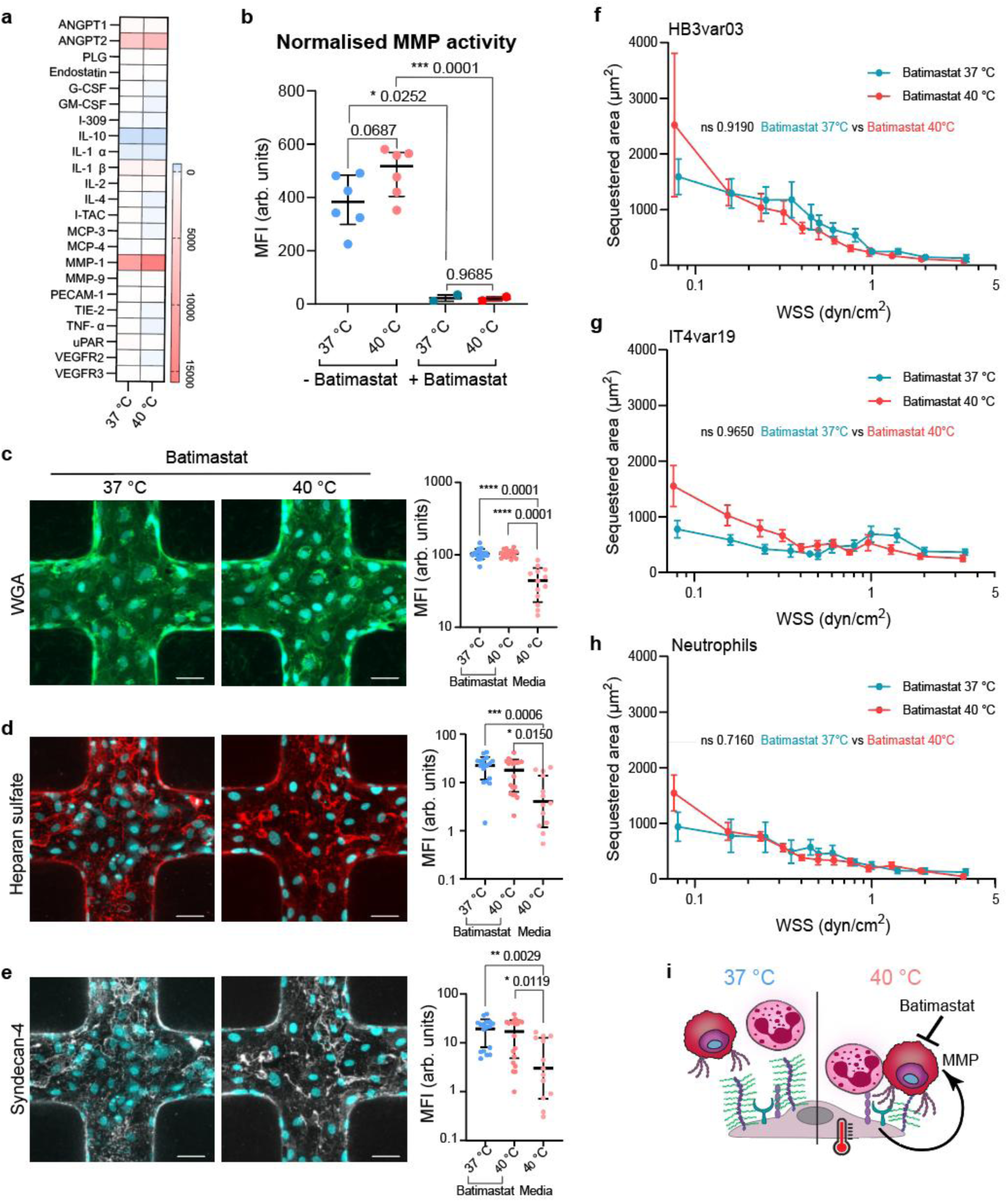
Analysis of glycocalyx-protective effect by batimastat, an MMP inhibitor. **a,** Quantification of angiogenesis-related molecule secretion from HBMEC monolayers after 1 h at 37 °C or 40 °C. Supernatants pooled from *n* =3 biological replicates and normalised as described in Methods. **b**, Quantification of MMP activity in supernatant taken from 2D HBMEC monolayers following 1-hour 37 °C or 40 °C treatment +/- batimastat (*n* = 6 for 1-hour 37 °C or 40 °C treatments and *n* = 2 for batimastat treatments). MMP activity was normalised to that of freshly-prepared endothelial media. Dot plots show mean ± s.d. where each dot represents independent biological replicates. Statistical analysis by the unpaired two-tailed Welch’s t-test. **c-e,** Left, representative z-projection immunofluorescence images of brain microvessels pretreated with batimastat and exposed to 37 °C or 40 °C for 1 h, labelled for WGA (**c**), heparan sulfate (**d**), syndecan-4 (**e**), and DAPI (cyan). Right, corresponding mean fluorescence intensity (MFI), normalised to cell number and compared with microvessels exposed to 40 °C (media). *n* = 4 brain microvessels per condition, with each dot representing an ROI (Dot plots show mean ± s.d.). Statistical analysis was performed using the Kruskal–Wallis with Dunn’s multiple-comparisons tests. Scale bars = 50 µm. **f-h,** HB3var03 (**f**), IT4var19 (**g**) and neutrophils (**h**) binding after batimastat-treatment of 3D brain microvessels at 37 °C or 40 °C (*n* = 4-7 brain microvessels). Mean ± error bars representing s.e.m. Statistical analysis between sequestered areas at 37 °C and 40 °C was determined using a linear mixed model to account for WSS-dependence. **i,** Proposed model summarising the mechanism by which febrile temperature enhances vascular adhesion.

## Discussion

Although fever generally exerts protective effects in the host during pathogenic diseases, our findings reveal that it may paradoxically amplify severe malaria pathology by promoting adhesion of circulating blood cells. Here we show that brief exposure to febrile temperature causes vascular glycocalyx shedding, increasing *P. falciparum*-iRBC and neutrophil adhesion. Enhanced cytoadhesion has important clinical implications: high parasite biomass in the microvasculature is a key component of severe malaria^44,45,13^, and together with neutrophil activation^46,34^ contributes to vascular obstruction and complications such as blood-brain barrier disruption and acute respiratory distress syndrome through inflammatory and vascular-disruptive pathways^47–49^. Consistent with our findings, clinical observations have linked glycocalyx breakdown to fatal outcomes^50^, and higher fever to adverse outcomes during cerebral malaria, including epilepsy, long-term disabilities, and cognitive impairments^51^. Together, the current study proposes that febrile temperatures amplify malaria-associated vascular pathology through endothelial glycocalyx degradation.

The glycocalyx is a dynamic network of proteoglycans and glycoproteins that forms a protective shield, limiting direct interaction of circulating blood cells with endothelial cells. Its breakdown is common in severe *P. falciparum*^50,52–54^, *vivax*, and *knowlesi* malaria^39^, while its recovery is slow, requiring about 20 hours after enzymatic degradation *in vitro*^55^ and up to 7 days *in vivo*^56^. Thus, the incomplete recovery we observed *in vitro* between consecutive febrile episodes is consistent with previous studies. The glycocalyx varies in thickness from 0.2 to 4.5 μm^57^, and major cytoadhesion receptors, such as ICAM-1 (∼19 nm)^58,59^ and the even smaller EPCR, remain largely concealed beneath this layer^31^. In contrast with neuraminidase treatment, febrile-range temperatures induced disruption and shedding of deeper components, such as syndecan-4 and heparan sulfate. This mechanistic distinction likely explains why neuraminidase treatment alone mostly recapitulates the full febrile phenotype at WSS lower than 1 dyn cm⁻². At higher WSS and flow velocities, iRBC–endothelial contact times are brief, making the temperature-associated extensive glycocalyx disruption critical for establishing stable adhesion. Thus, our results indicate that febrile temperature enhances cytoadhesion not by upregulating ICAM-1 or EPCR expression, but by increasing the accessibility of these otherwise shielded receptors.

Importantly, glycocalyx disruption was not restricted to the brain microvasculature. Pulmonary microvessels also exhibited glycocalyx shedding following febrile exposure, although to a lesser extent, suggesting that hyperthermia induces a conserved endothelial response with organ-specific susceptibility. These differences may reflect endothelial organotypic specialisation, as the brain endothelium possesses a thicker glycocalyx than that of the pulmonary endothelium^60^. Our microvessel models reproduced these organotypic differences, with pulmonary microvessels exhibiting lower basal sialic acid and N-acetylglucosamine at 37 °C. A less developed basal pulmonary glycocalyx may limit the absolute extent of glycocalyx loss following febrile exposure providing a plausible explanation for the smaller increase in iRBC adhesion in the lung model. Conversely, we propose that the brain’s more extensive baseline glycocalyx provides a greater receptor shielding effect, which may explain its greater susceptibility to fever-induced increases in receptor accessibility. Consistent with this mechanism, post-mortem analyses of CM patients reveal distinct glycocalyx shedding within the brain microvasculature, confirming that this process occurs *in vivo*^61^. Together, these observations suggest that while fever-mediated glycocalyx breakdown is a common mechanism across distinct vascular beds, differences in glycocalyx architecture may contribute to organ-specific vascular responses to fever and represent an underappreciated determinant of parasite tropism during severe malaria.

Our study also provides insights into how febrile temperatures rapidly disrupt the glycocalyx. The short temperature increase did not induce major changes at the protein level. Instead, it triggered rapid release of active MMP-1 and angiopoietin-2, likely from intracellular endothelial vesicles^62^, and increased in MMP1 activity. Consistent with this mechanism, studies in severe malaria patients have reported increased MMP expression, serum secretion, and localisation to sites of vascular damage^63,64^, and studies in stroke have shown that hyperthermia upregulates MMP activity, contributing to basement membrane breakdown^65^. In contrast, heparanase levels were unchanged following febrile exposure, arguing against a major contribution in our model. MMP inhibition with batimastat preserved the glycocalyx and prevented temperature-dependent increases in both iRBC and neutrophil adhesion, supporting MMP activity as the principal driver of endothelial glycocalyx dysfunction. This is in agreement with experimental CM studies showing delayed mortality following MMP inhibition and glycocalyx preservation^66^. Although this suggests the possibility of its use as an adjunctive therapy, batimastat presents broad-spectrum MMP inhibition, lending to off-target effects that limit its clinical development and encoraurages the development of more specific MMP inhibitors^67^. Despite this limitation, our findings further underscore the need for more aggressive fever management during severe malaria treatment^9^.

Beyond promoting parasite sequestration, febrile temperature also amplifies host immune interactions with the vasculature. Previous studies have shown that hyperthermia accelerates neutrophil migration^68^, extracellular trap formation^69^, and enhances leukocyte adhesion^40,70–73^. Similarly, we found that febrile temperature increased neutrophil adhesion through ICAM-1, given the limited E-selectin expression in our endothelial model. Neutrophil recruitment initiates through selectin-mediated interactions followed by engagement of endothelial ICAM-1, which mediates firm adhesion and subsequent transmigration. Although selectin ligands are concentrated on neutrophil microvilli, which may penetrate the glycocalyx, ICAM-binding integrins are broadly distributed across the neutrophil cell body^74^. This differential membrane spatial organisation may explain why glycocalyx shedding preferentially enhanced ICAM-1-dependent neutrophil adhesion in our model. The greater enhancement of iRBC adhesion compared with neutrophils may also reflect differences in their mechanical properties, as iRBCs are more rigid whereas neutrophils are highly deformable and dynamically remodel their shape during adhesion^75^. Nevertheless, the fact that glycocalyx degradation enhances the adhesion of both iRBC and neutrophils, despite differences in adhesion receptors and mechanics, suggest that fever-induced glycocalyx remodelling broadly increases endothelial adhesiveness. Only very few neutrophils were observed outside the vessels, indicating that transmigration was not substantially detected following increased adhesion. Future studies incorporating chemokine gradients or prolonged neutrophil incubation times at febrile temperatures could better determine whether temperature contributes to transmigration and if sustained neutrophil-endothelial interactions contribute to vascular inflammation and barrier dysfunction.

The 3D flow-based microvessel chip overcomes key limitations of traditional assays for studying febrile-temperature effects on *P. falciparum*-iRBC interactions. By preserving the native endothelial glycocalyx, the receptor mobility, and vascular flow, this model captures features that are absent from assays using recombinant proteins. This distinction may account for previously reported discrepancies, where heating reduced binding to recombinant receptors in force-spectroscopy assays^17,76^ but increased cytoadhesion of iRBCs to endothelial cells^17^. The 3D platform has been previously validated in studies of *P. falciparum* interactions with EPCR and ICAM-1^77^, and in testing inhibitory effects of plasma^78^ or monoclonal antibodies^29^. An advantage of our system is that it allows decoupling of the temperature-effect of fever, independently of fever-associated pro-inflammatory cytokines that upregulate endothelial receptors, such as ICAM-1. The independent tuning of physical and biological parameters also represents an advantage over hyperthermia animal models. A limitation of the study is that rapid mechanical modifications of *P. falciparum-*iRBC or neutrophils upon temperature exposure could further contribute to increased cytoadhesion. However, the strong inhibition of binding with batimastat suggests that glycocalyx disruption is a major contributor to the increased binding. Future studies should examine the impact of glycocalyx impairment on bond dynamics under flow, the combined effect of hyperthermia and inflammatory signalling on cell adhesion, and extend the model with additional blood–brain barrier cell types or immune cells to directly assess febrile contributions towards CM pathology. Our study has implications beyond malaria. Glycocalyx dysregulation is increasingly recognised as a central determinant of vascular vulnerability, implicated in viral lung injury^32,79^, sepsis^80^, ageing and neurodegeneration^81^. Consistently, hyperthermia worsens outcomes in stroke^8^ and hypoxic–ischemic brain injury^82^ by disrupting the blood–brain barrier and causing brain aedema^83^ —paralleling CM pathology. Our findings therefore extend to a broader principle: febrile temperature destabilises the glycocalyx and amplifies microvascular pathology across infectious and inflammatory diseases.

## Methods

### Body temperature measurements in patients

Body temperature data were collected from 146 children (ages 2–11) who received either standard-of-care antipyretic therapy, following WHO recommendations, with acetaminophen (15 mg/kg every 6 hours as needed for temperature ≥ 38.5°C), or aggressive antipyretic therapy with prophylactic acetaminophen (30 mg/kg loading dose, followed by 15 mg/kg every 6 hours) plus ibuprofen (10 mg/kg every 6 hours). Clinical temperature recordings were analysed to identify febrile episodes reaching ≥ 40 °C. The duration of individual febrile episodes was also quantified from the clinical recordings and used to reproduce clinically relevant short febrile exposures (1 h at 40 °C) in the microvascular models. In patients presenting at least two consecutive febrile peaks (n = 17), the time interval between the first and second peaks was calculated, and the mean interval was used to design the cyclic fever regimen applied *in vitro*. Participants were enrolled at Queen Elizabeth Central Hospital (Blantyre, Malawi)^6,9^. All had *P. falciparum* infection confirmed by blood smear or rapid diagnostic test and presented with central nervous system symptoms (impaired consciousness and/or seizures) of whom 72–83% met CM criteria. Temperatures were recorded via a body patch every 2–4 minutes over a 72-hour period, during which most children defervesce. Ethics approvals: College of Medicine Research and Ethics Committee (Malawi): P.10/17/2298.

### Primary cell culture and 3D microvessel fabrication

Primary human brain microvascular endothelial cells (HBMECs; Cell Systems, ACBRI 376) and primary human pulmonary artery endothelial cells (HPAECs; Lonza, CC-2530) were cultured in poly-L-lysine-coated (Sigma, P8920) T75 flasks at 37 °C and 5% CO₂. HBMECs were maintained in Endothelial Growth Medium-2MV (EGM-2MV; Lonza, CC-3202) supplemented with 5% fetal bovine serum, and HPAECs in VascuLife® VEGF Endothelial Medium (Lifeline Cell Technology, LL-0003) supplemented with 2% fetal bovine serum, according to the manufacturers’ instructions. All experiments were performed using mycoplasma-free cells between passages 7 and 9. 3D microvessels were fabricated as described previously^77^. Briefly, the upper microvascular network was generated by casting type I collagen (7.5 mg/ml) between a patterned polydimethylsiloxane (PDMS) mold containing a 13 × 13 vascular grid and a PMMA jig. The lower compartment consisted of a flat collagen layer compressed between a flat PDMS stamp and a 22 × 22 mm glass coverslip. After collagen polymerization for 30 min at 37 °C, the PDMS stamps were removed and the two components assembled to generate a perfusable 3D collagen microvascular network. HBMECs or HPAECs were seeded at 7 × 10⁶ cells/ml under gravity-driven flow by sequential addition of 8 µl aliquots to the device inlet until complete endothelialisation of the network was achieved. Microvessels were maintained in the corresponding culture medium for 3 days and perfused every 12 h by gravity-driven flow before *P. falciparum* adhesion assays. To mimic cyclic fever observed in patients, after 3 days brain microvessels were exposed to one or two 1-h incubations at either 37 °C or 40 °C, separated by a 6-h recovery period at 37 °C. Microvessels were then fixed in 4% PFA for 20 min, washed twice with PBS, and processed for immunofluorescence staining and imaging.

### Numerical simulation of wall shear stress rates

The flow profile within the 3D brain microvessels was simulated using COMSOL Multiphysics software, as previously described^77^. Flow in the microvessel network (diameter 120 µm) was assumed to be laminar, and the stationary solver for laminar flow was used with predefined Navier-Stokes equations. Due to the low hematocrit (<0.1%) used during perfusion, flow was assumed to be Newtonian, and WSS rates were calculated based on fluid viscosity of water or culture medium at 37 °C (viscosity of 6.922 × 10^-4^ Pa s and density of 993.3 kg/m^3^) and 40 °C (viscosity of 6.539 × 10^-4^ Pa s and density of 992.2 kg/m^3^). The inlet boundary conditions were defined for the perfusion flow rate of 10 μl/min, and the outlet boundary conditions were set at zero pressure.

### Particle tracking and particle image velocimetry (PIV)

Flow velocity was determined by tracking 1.0-µm fluorescent yellow-green carboxylate-modified polystyrene latex beads (L4655, Sigma-Aldrich) suspended in endothelial culture medium as they flowed through the centerline of the microvessels. Time-lapse videos were acquired on a Zeiss LSM 900 microscope at 120 frames/s for 10 s at both 37 °C and 40 °C from six regions of the microvascular network corresponding to low-, intermediate-, and high-shear locations along opposite vessel edges. Image sequences were background-subtracted in Fiji before particle trajectories were quantified using the TrackMate plugin^84^. PIV was performed in MATLAB using PIVlab3.00^85^. Following background subtraction and filtering by pixel size, velocity vector fields were computed using a multi-pass cross-correlation algorithm with interrogation windows of 256, 128, and 64 pixels and step sizes of 64, 64, and 32 pixels, respectively, and velocity profiles were extracted along the microvessel channels.

### Plasmodium falciparum culture

*P. falciparum* lines HB3var03, IT4var19, and IT4var31 (kindly provided by Joe Smith’s lab, Seattle Children’s Research Institute) and NF54var2csa (kindly provided by Moritz Treeck, Gulbenkian Institute for Molecular Medicine) were cultured in human O+ RBCs obtained by the Banc de Sang i Teixits, Barcelona, in RPMI-1640 medium (Gibco) containing 25 mM HEPES, 1.5 g/L gentamicin, 0.4 mM hypoxanthine, 26.8 mM sodium bicarbonate, 5 mM glucose, and 10% human type AB+ serum. Parasites were grown in a gas mixture of 90% N_2_, 5% CO_2_, and 5% O_2_ and parasitemia and daily checked by Giemsa staining. Cultures were regularly panned and monitored for correct PfEMP1 expression on primary HBMECs. *P. falciparum* parasites were synchronized weekly using 5% sorbitol to select for ring-stage parasites. The var gene transcription profile of ring-stage iRBCs was analysed by qRT-PCR with IT4, HB3, and 3D7 (NF54 strain is identical to the NF54-derived clone 3D7) *var* strain-specific primer sets. Real-time PCR reactions were performed on a Roche LightCycler 480 II using Power SYBR Green Master Mix and previously published conditions for the respective dataset (Extended Data Fig. 3). Transcript levels were normalized to the housekeeping gene *SARS* (seryl-tRNA synthetase).

### Temperature-dependent iRBC and neutrophil binding assays in 3D microvessels

Immediately before experiments, *P. falciparum* cultures were enriched for mature-stage infected erythrocytes (iRBCs) using a MACS cell separator with LD columns (Miltenyi Biotec, 130-042-901) and diluted to 5 × 10⁶ cells/ml in the corresponding endothelial cell culture medium. iRBCs were labeled with PKH26 Red Fluorescent Cell Linker Midi Kit (Sigma, MIDI26-1KT) according to the manufacturer’s instructions. For neutrophil binding experiments, neutrophils were isolated from whole blood samples obtained from the Catalan Blood and Tissue Bank using the EasySep Direct Human Neutrophil Isolation Kit (StemCell Technologies, 19666) according to the manufacturer’s instructions. Isolated neutrophils were labeled with PKH26 and resuspended at 2 × 10⁶ neutrophils/ml in RPMI 1640 (Sigma) supplemented with 5% heat-inactivated fetal bovine serum (Gibco), 0.1% vascular endothelial growth factor (Lonza), 0.4% recombinant human FGF (Lonza), 0.1% ascorbic acid (Lonza), 0.1% R3-insulin-like growth factor (Lonza), 0. 1% recombinant human epithelial growth factor (Lonza), 0.5% penicillin/streptomycin (Gibco) and 1% Glutamax (Gibco). 3D microvessels (both HBMECs and HPAECs) were pre-incubated for 1 h at either 37 °C or 40 °C. Subsequently, iRBCs or neutrophils maintained at 37 °C were perfused through the microvessels for 30 min either at 37 °C or 40 °C condition, at a flow rate of 10 µl/min using a syringe infusion pump (KD Scientific, KDS220). Following perfusion, microvessels were washed for 5 min with EGM-2MV or VascuLife® endothelial medium for brain and lung microvessels, respectively, at the same flow rate. Samples were fixed with 4% paraformaldehyde for 20 min, washed twice with PBS for 10 min, and stained with DAPI. For receptor-blocking experiments, microvessels were pre-incubated for 20 min at the experimental temperature with blocking antibodies against endothelial ICAM-1 (mouse anti-human ICAM-1 mAb 15.2; Abcam, ab20; 5 µg/ml) or EPCR (rat anti-human EPCR mAb 252; Sigma-Aldrich, E6280-200UL; 50 µg/ml), together with the corresponding mouse or rat IgG isotype controls. For inhibition of PfEMP1 CIDRα1-mediated binding, iRBCs were incubated for 30 min at room temperature with either human IgG1 isotype control (BioLegend, 403502; 0.25 mg/ml) or the CIDRα1-blocking monoclonal antibody C7 (0.25 mg ml⁻¹) before perfusion, as previously demonstrated^29^. For MMP inhibition experiments, microvessels were treated with the MMP inhibitor batimastat (R&D Systems, 2961) at 5 µM for 2 h prior the temperature exposure and binding assays. Each microvessel device was used once per experimental condition.

### Binding quantification

For each microvessel device, images were acquired from the edge regions of the 13 × 13 microvascular network using a Zeiss LSM 980 Airyscan 2 microscope equipped with a 10× objective (NA 0.3). Cytoadhered iRBCs or neutrophils labeled with PKH26 membrane dye were detected using a 555 nm laser, while DAPI-stained parasite and HBMEC nuclei using a 405 nm laser. DAPI staining was used to confirm uniform endothelial coverage throughout the microvascular network. Images were acquired with a 3 µm z-step, and maximum-intensity projections of the vessel bottom were generated from z-stacks using Fiji (ImageJ v1.52b) software. The cytoadhered area was quantified as the area occupied by attached iRBCs or neutrophils within 12 predefined regions along the vessel edges, as previously described^29^. To minimise flow-related variability, quantification was restricted to central channel regions with fully developed laminar flow, excluding branch junctions and inlet/outlet regions. Each microvessel device was considered an independent biological replicate. Up to four vessel edges per device were analysed, with 12 predefined WSS regions quantified within these edges (see Expanded Data Fig.1 for a graphical schematic).

### Immunofluorescence microscopy of 3D microvessels

Fixed 3D microvessels were incubated with Background Buster (Innovex, NB306; 30 min) and blocked with 2% BSA and 0.1% Triton X-100 in PBS for 1 h. Samples were incubated overnight at 4 °C with primary antibodies against VE-cadherin (Abcam, ab33168; 1:100), vWF (Bio-Rad, AHP062; 1:100), Syndecan-4 (R&D Systems, AF2918; 1:50), or heparan sulfate (Amsbio, 370255-S; 1:100). After washing, samples were incubated with Alexa Fluor-conjugated secondary antibodies (Invitrogen; 1:250) and DAPI (8 µg/ml) for 1 h at room temperature. Glycocalyx sialic acids were visualised using FITC-WGA (VWR, 29022-1; 1:50) or fluorescein-conjugated *Sambucus nigra* lectin (Vector Laboratories; 1:100) for 20 min. Images were acquired using a Zeiss LSM 980 Airyscan 2 confocal microscope (20×/0.8 NA objective; 1-µm z-steps). Three-dimensional reconstructions were generated using ZEN v3.3 (Zeiss) and Arivis Vision4D v3.5.1.

### Cytokine/angiogenesis antibody array and ELISA assays

Supernatants from HBMEC monolayers cultured in 6-well plates (*n* =3 biological replicates) were analysed using a Human Angiogenesis Antibody Array (Abcam, ab193655) according to the manufacturer’s instructions. Chemiluminescence signals were background-corrected using the negative controls, normalised to the positive controls, and compared between samples exposed to 37 °C and 40 °C for 1 h. Blots were imaged (1 min exposure) using a Fusion FX Spectra system (Vilber, France). For ELISA assays, media from the outlet reservoir of each device was collected after incubation for 1 h at 37 °C and 40 °C from *n* = 3-6 experiments at day 3. Quantification of Syndecan-4, Syndecan-1, and Heparanase was performed using the Human Syndecan-4 DuoSet ELISA kit (R&D Systems, DY2918), Human Syndecan-1 DuoSet ELISA kit (R&D Systems, DY2780), and Human Heparanase ELISA kit (Abcam, ab256401), according to manufacturer’s instructions.

### Proteomics

<u>Sample Preparation:</u> HBMEC cells were cultured on flasks for three to four days until confluent, and then incubated for 1 h either at 37 °C or 40 °C (total of 3 biological replicates per condition). Cells were lysed in cold RIPA buffer with 1× protease inhibitors, incubated on ice for 15 min, sonicated (5 pulses of 5 min at 0/0.5 high, 5°C), and centrifuged at ∼14,000 × g for 15 min. Protein concentration was determined using the BCA assay by incubating 20 µl of samples with 200 µl of BCA working reagent (50:1, Reagent A:B) at 37°C for 30 min, followed by absorbance measurement at 562 nm. Samples were subjected to the SP3 protocol^86,87^ on a King Fisher Apex Platform (Thermo Fisher) and peptides were eluted by tryptic digestion (sequencing grade trypsin, Promega) for 5 h at 37°C. Peptides were labelled with TMT6plex^88^ Isobaric Label Reagent (ThermoFisher) according the manufacturer’s instructions. In short, 0.8 mg reagent was dissolved in 42 µl acetonitrile (100%) and 4 µl of stock was added and incubated for 1h room temperature. Followed by quenching the reaction with 5% hydroxylamine for 15 min at RT. Samples were combined and for further sample clean up an OASIS® HLB µElution Plate (Waters) was used. The TMT6-labelled proteome was fractionated by high-pH reversed-phase carried out on an Agilent 1200 Infinity high-performance liquid chromatography system, equipped with a Gemini C18 column (3 μm, 110 Å, 100 x 1.0 mm, Phenomenex). 48 fractions were collected along with the LC separation that were subsequently pooled into 12 fractions. Pooled fractions were dried under vacuum centrifugation and reconstituted in 10 μl 1% formic acid, 4% acetonitrile and then stored at -80 °C until LC-MS analysis. <u>Data acquisition.</u> Samples were measured on an Q Exactive™ Mass Spectrometer (Thermo) coupled to an UltiMate 3000 RSLC nano LC system (Dionex). Sample was concentrated on a C18 µ-Precolumn (Acclaim PepMap 100, 5µm, 300 µm i.d. x 5 mm, 100 Å) and resolved on a nanoEase™ M/Z HSS T3 column from Waters (75 µm x 250 mm C18, 1.8 µm, 100 Å). Trapping was carried out at a constant flowrate of 30 µl/min 0.5% trifluoroacetic acid in water for 4 min. Subsequently, peptides were eluted via the analytical column (solvent A: 0.1% formic acid in water, 3% DMSO) with a constant flow of 0.3 µl/min, with increasing percentage of solvent B (0.1% formic acid in acetonitrile, 3% DMSO) from 2% to 8% in 6 min, 8% to 28% in 37 min, from 28% to 40% in 9 min, followed by an increase of B from 40-80% for 3 min and a re-equilibration back to 2% B for 5 min. The peptides were introduced into the mass spectrometer via a Pico-Tip Emitter 360 µm OD x 20 µm ID; 10 µm tip (New Objective) and an applied spray voltage of 2.4 kV. The capillary temperature was at 275°C. Settings for the Q Exactive were: Full mass scan (MS1) was acquired with mass range 375-1200 m/z, profile mode, in the orbitrap, resolution of 70000, fill time 10 ms. AGC target 3E6. Data dependent acquisition (DDA) was performed with the resolution of the Orbitrap set to 17500, fill time 50 ms, AGC target of 9E1 ions. Normalized collision energy of 32, HCD, fixed first mass 110 m/z. <u>Database</u> <u>search</u>. Fragpipe v21.1 with MSFragger v4.0^89^ was used to process the acquired data, which was searched against the homo sapiens Uniprot proteome database (UP000005640, ID9606, 20594 entries, release October 2022) with common contaminants and reversed sequences included. The following modifications were considered as fixed modification: Carbamidomethyl (C) and TMT6 (K). As variable modifications: Acetyl (Protein N-term), Oxidation (M) and TMT6 (N-term). For the MS1 and MS2 scans a mass error tolerance of 20 ppm was set. Further parameters were: Trypsin as protease with an allowance of maximum two missed cleavages; Minimum peptide length of seven amino acids; The false discovery rate on peptide and protein level was set to 0.01. <u>Data analysis.</u> The raw output files of FragPipe (protein.tsv files) were processed using the R programming environment (ISBN 3-900051-07-0). Initial data processing included filtering out contaminants and reverse proteins. Only proteins quantified with at least 2 razor peptides (with Razor.Peptides >= 2) were considered for further analysis. 4203 proteins passed the quality control filters. In order to correct for technical variability, batch effects were removed using the ‘removeBatchEffect’ function of the limma package on the log2 transformed raw TMT reporter ion intensities (’channel’ columns)^90^. Subsequently, normalization was performed using the ‘normalizeVSN’ function of the limma package (VSN - variance stabilization normalization^91^). Differential expression analysis was performed using the moderated t-test provided by the limma package^90^. The model accounted for replicate information by including it as a factor in the design matrix passed to the ‘lmFit’ function. Proteins were annotated as hits if they had a false discovery rate (FDR) below 0.05 and an absolute fold change greater than 2. Proteins were considered candidates if they had an FDR below 0.2 and an absolute fold change greater than 1.5.

### Western blot

10^5^ HBMECs/well were seeded into 6-well plates and grown for 3 days. Upon formation of a confluent monolayer on day 3, cells were incubated at 37 or 40°C for 1, 6 or 24 h. Cells were then washed with 4°C PBS and lysed for 15 minutes with 4°C Pierce RIPA buffer (Invitrogen, 89900) and 1x Protease inhibitor (Invitrogen, A32963). The cellular components were suspended using a cell scrapper and the resulting cell lysate solution was spun at 18000 x g, 4°C to remove cell debris. Quantification of protein concentrations were performed using a BCA assay (Invitrogen, 23227) following manufacturer’s instructions. 20 mg of protein or 5 μl of Precision Plus Protein WesternC Blotting Standard (Bio-Rad, 1610376) was ran on precast Mini-PROTEAN TGX gels (Bio-Rad, 4561034) at 170 V for 1 h. Gels were then transferred to nitrocellulose membranes (Bio-Rad, 1704159) using a Trans-blot Turbo transfer machine programmed to mixed molecular weight (2.5 A, 25 V). Membranes were incubated for 30 seconds in 100% methanol and blocked in 5% BSA-tris-buffered saline (TBS) for 2 h on a shaker set to 50 rpm. Blocked membranes were incubated with 1:1000 rabbit anti-heparanase (Cell Signaling, 99756S), 1:1000 rabbit anti-prostaglandin E synthase 2 (Thermofisher, 16817563), 1:1000 rabbit anti-heat shock protein 70 (R&D Systems, AF1663-SP) or 1:1000 mouse anti-β-tubulin (Invitrogen, MA5-16308) in 3% BSA-TBS+0.2% Tween-20 overnight at 4°C on a shaker. Membranes were then washed 3 times for 5 minutes with TBS+0.2% Tween-20 and incubated for 1 h at RT on a shaker with 1:10000 IRDye goat anti-mouse 680R (LI-COR Biotech, 926-68070) and 1:20000 IRDye goat anti-rabbit 800CW (LI-COR Biotech, 926-32211) in 3% BSA-TBS+0.2% Tween-20+0.01% sodium dodecyl sulphate. After being washed 4 times for 5 minutes with TBS+0.2% Tween-20, membranes were rinsed in dH_2_O and imaged using a LI-COR Odyssey M scanner. Acquired images underwent densitometry analysis using Fiji where a ROI was manually fitted to each band at the correct molecular weight (50 kDa for mature heparinase, 30 kDa for prostaglandin E synthase 2, 70 kDa for heat shock protein 70 and 55 kDa for β-tubulin), including a ROI of equal size on an area of the membrane with no bands to be used for background subtraction. Integrated density values of each protein band were subtracted by the background integrated density value and then normalized to that of the β-tubulin control by division.

### Flow cytometry

HBMECs and HPAECs were cultured to confluence in T75 flasks, transferred to 6-well plates, and incubated at 37 °C or 40 °C for 1 h. Cells were rinsed twice with 1x HBSS, detached using 15 mM EDTA in HBSS followed by a PBS wash containing 0.5% BSA at 1000 rpm for 5 mins at room temperature. 1 × 10⁵ cells per well were transferred to 96-well plates and stained with the indicated antibodies on ice for 30 min per staining step, with washes between incubations, before analysis on a BD LSRFortessa flow cytometer. The following antibodies were used: goat anti-human EPCR (5 μg/ml; R&D Systems, AF2245), followed by donkey anti-goat Alexa Fluor 647-conjugated secondary antibody (1:400; Invitrogen, A32849); PE-conjugated mouse anti-human E-selectin (5 μl/test; Thermofisher, 12-0627-42); PE-conjugated anti-human ICAM-1 (2 μl per 10⁶ cells; Miltenyi Biotec, REA266, 130-120-711) and relative isotype control PE-conjugated human IgG1 (Miltenyi, 130-113-471); FITC-conjugated anti-human CD31 (5 μl per 10⁵ cells; BD Pharmingen, 560984) and relative FITC-labelled mouse IgG1 (Abcam, ab18447); FITC-labelled anti-human CD36 (10 μl per 10⁵ cells; Abcam, ab39022) and relative FITC-conjugated mouse IgG1 isotype control (2 μg/ml; Abcam, ab106163). All experiments were performed on live cells, and dead cells were excluded using DAPI (ThermoFisher, D21490; 1:600 dilution). Data were analysed using FlowJo v10 software (Tree Star Inc.). Debris, doublets and non-viable cells were excluded from each cell population (**Extended Data Figs. 7c)**. Then, the percentage of positive gated cells and mean fluorescence intensity (MFI) were quantified for CD31, CD36, E-selectin, EPCR and ICAM-1 from the live cell gate. The gates were determined by using marker-negative cells of each cell population (**Extended Data Figs. 7c)**. Co-expression of ICAM-1 and EPCR was assessed using bivariate plots for classification of the endothelial cells into ICAM-1⁻EPCR⁻, ICAM-1⁺EPCR⁻, ICAM-1⁻EPCR⁺, and ICAM-1⁺EPCR⁺ subsets. ICAM-1⁺EPCR⁺ subset was described as the population co-expressing ICAM-1 and EPCR on the same cell.

### In vitro matrix metalloproteinase assay

10^5^ HBMECs/well were seeded into 6-well plates and grown for 3 days. Upon formation of a confluent monolayer on day 3, media was changed and replaced with 500 μl of EGM-2MV to concentrate secreted MMPs. HBMECs were incubated at 37 or 40°C for 1 h, +/- 2 h preincubation with 5 μM Batimastat, and supernatant was collected. 1:100 p-Aminophenylmercuric acetate (APMA) (Sigma, 164610-700MG) was added to supernatant from each condition, including an EGM-2MV stock control that was not cultured with cells, and incubated at 37°C for 1 h. Each supernatant was then added in 100 μL duplicates to a clear bottom 96 well plate and incubated +/- 1 μM MMP Mca-KPLGL-Dpa-AR-NH2 fluorogenic peptide substrate (R&D Systems, ES010) at 37°C for 1 h on a shaker set to 100 rpm in the dark. Resulting fluorescence given by peptide cleavage was measured using a Tecan Infinite M1000 Pro microplate reader using an excitation wavelength of 320 nm and emission wavelength of 405 nm. Fluorescence intensity values were first normalised by subtracting the fluorescence values of the EGM-2MV stock + APMA controls lacking the fluorogenic peptide substrate, to remove the non-specific fluorescence background. The resulting values were then normalised a second time by subtracting the fluorescence values of the EGM-2MV stock + APMA controls containing the fluorogenic peptide substrate to account for any intrinsic MMP activity present in the stock medium.

### Statistical analysis

Binding assays in 3D microvessels were analysed using linear mixed-effects models (LMM) (lme4 package - Rstudio) in the log-transformed sequestered area (log(area + 1)). Experimental condition and, where applicable, WSS group were included as fixed effects, whereas device and edge nested within device were included as random intercepts. Degrees of freedom were estimated using the Kenward–Roger approximation. Model residuals were assessed by Q–Q plots. Pairwise comparisons (p -values) were performed using estimated marginal means with Tukey adjustment for multiple testing. We performed a continuous sensitivity analysis by systematically varying the WSS threshold from the 20th to the 100th percentile of the pooled WSS distribution in 20% increments and refitting the LMM at each threshold. For all other statistical analyses (e.g. boxplots), GraphPad Prism (v10.2.0) was used. Normality was assessed using the Shapiro–Wilk test, and variance homogeneity was assessed using Levene’s test. For two-group comparisons, paired Wilcoxon signed-rank tests were used for paired data, and unpaired Welch’s *t*-tests or Mann–Whitney *U* tests were used for normally or non-normally distributed data, respectively. For comparisons involving more than two groups, normally distributed data with homogeneous variances were analysed using one-way ANOVA with Tukey’s multiple-comparisons test and non-normally distributed data were analysed using the Kruskal–Wallis test with Dunn’s multiple-comparisons test. Statistical significance was defined as *p < 0.05, **p < 0.01, ***p < 0.001.

## Data availability

The mass spectrometry proteomics data have been deposited to the ProteomeXchange Consortium via the PRIDE [1] partner repository with the dataset identifier PXD063878.

## Acknowledgements

The authors thank Jennifer Schwarz (Proteomics Core Facility, EMBL Heidelberg); Felix Schneider and Sarah Kaspar (Data Science Centre, EMBL Heidelberg) for their support with the linear mixed model statistical analysis; the CRG-UPF Flow Cytometry Facility; the Microscopy Core Facility at the Max-Planck-Zentrum für Physik und Medizin; Alina Batzilla for experimental advice, and Moritz Treeck from the Gulbenkian Institute for Molecular Medicine for providing the NF54var2csa parasite line.

## Authorship contributions

Conceptualization: VI, MB; Methodology & validation: VI, RKML, MB; Data collection: VI, RKML, ORO, SSS, LH, MGM, IRA, GMH, BLG, KBS, GLB; Formal analysis: VI, RKML, FS, ORO, MGM; Visualization: VI, MB; Writing original draft: VI, RKML, MB; Supervision: MB; Project Administration: VI, MB; Funding Acquisition: VI, MB; All authors gave final approval for publication.

