## Supplementary Data for "Febrile temperature enhances *Plasmodium falciparum* and neutrophil adhesion by disrupting the endothelial glycocalyx"

### Extended Data

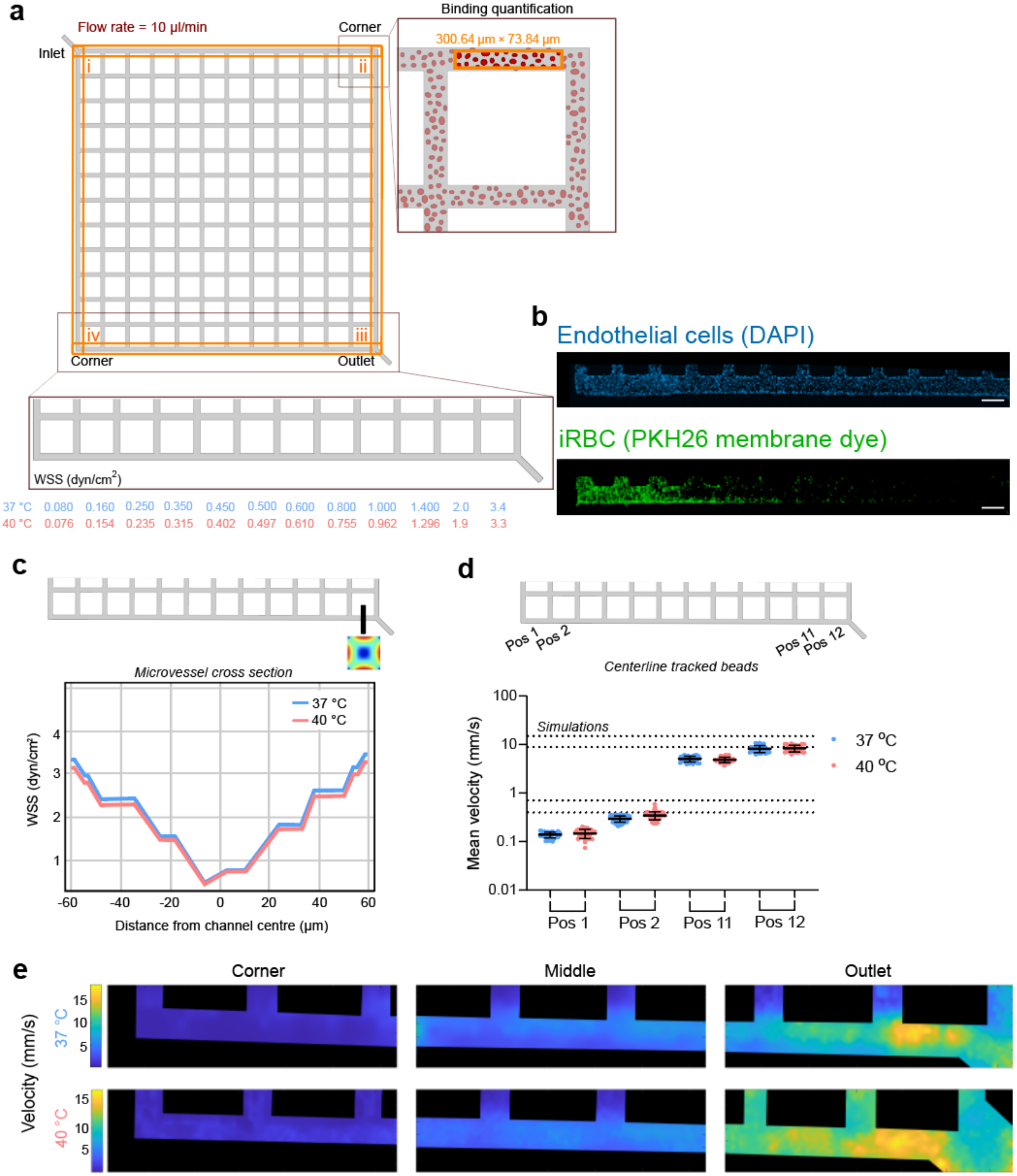


**Extended Data Figure 1. Quantification of iRBC binding and characterisation of flow conditions across the microvessel network.**

**a,** Schematic illustrating the workflow used to quantify *Plasmodium falciparum*-infected red blood cell (iRBC) sequestration within the 13x13 microvessel network. Adhesion was analysed along the four microvessel edges corresponding to the inlet (i), outlet (iii), and two opposite corner regions (ii and iv). The orange rectangle indicates the region of interest (300.84 × 73.84 $\mu m^2$) used for binding quantification along each edge. Bound iRBCs were quantified as the sequestered area within the region of interest and related to the local wall shear stress (WSS; dyn/cm$^2$). Simulated WSS values for each edge are shown for both 37 °C and 40 °C under a constant inlet flow rate of 10 µl/min.

**b,** Representative immunofluorescence images showing endothelial coverage and iRBC sequestration within a microvessel. Two-dimensional maximum-intensity projections of z-stack images are shown. Endothelial cell nuclei were stained with DAPI (blue), and iRBCs were labelled with PKH26 membrane dye (green). Scale bars, 100 µm.

**c,** Simulated cross-sectional WSS profile across the first branch of the microvessel network. The black line in the schematic indicates the cross-section used for the WSS profile. WSS distributions were calculated using COMSOL Multiphysics based on the viscosity and density of the perfusion medium at 37 °C and 40 °C before endothelial cell seeding (details in Methods).

**d,** Experimental validation of flow velocity by tracking 1-µm fluorescent beads along the centreline of representative microvessels at 37 °C and 40 °C. Measurements were performed at the two lowest WSS positions (Pos 1 and Pos 2) and at the two highest WSS positions (Pos 11 and Pos 12). Dotted lines indicate the corresponding centreline velocities predicted by COMSOL simulations before endothelial cell addition. Each dot represents an individual tracked bead (~40 per position). Dot plots show mean ± s.d. Statistical analysis was performed using the Mann–Whitney *U* test.

**e,** Representative particle image velocimetry (PIV) velocity maps acquired at the corner (low WSS), middle region, and outlet (high WSS) of representative microvessels at 37 °C and 40 °C, confirming the spatial distribution of flow velocities across the network.

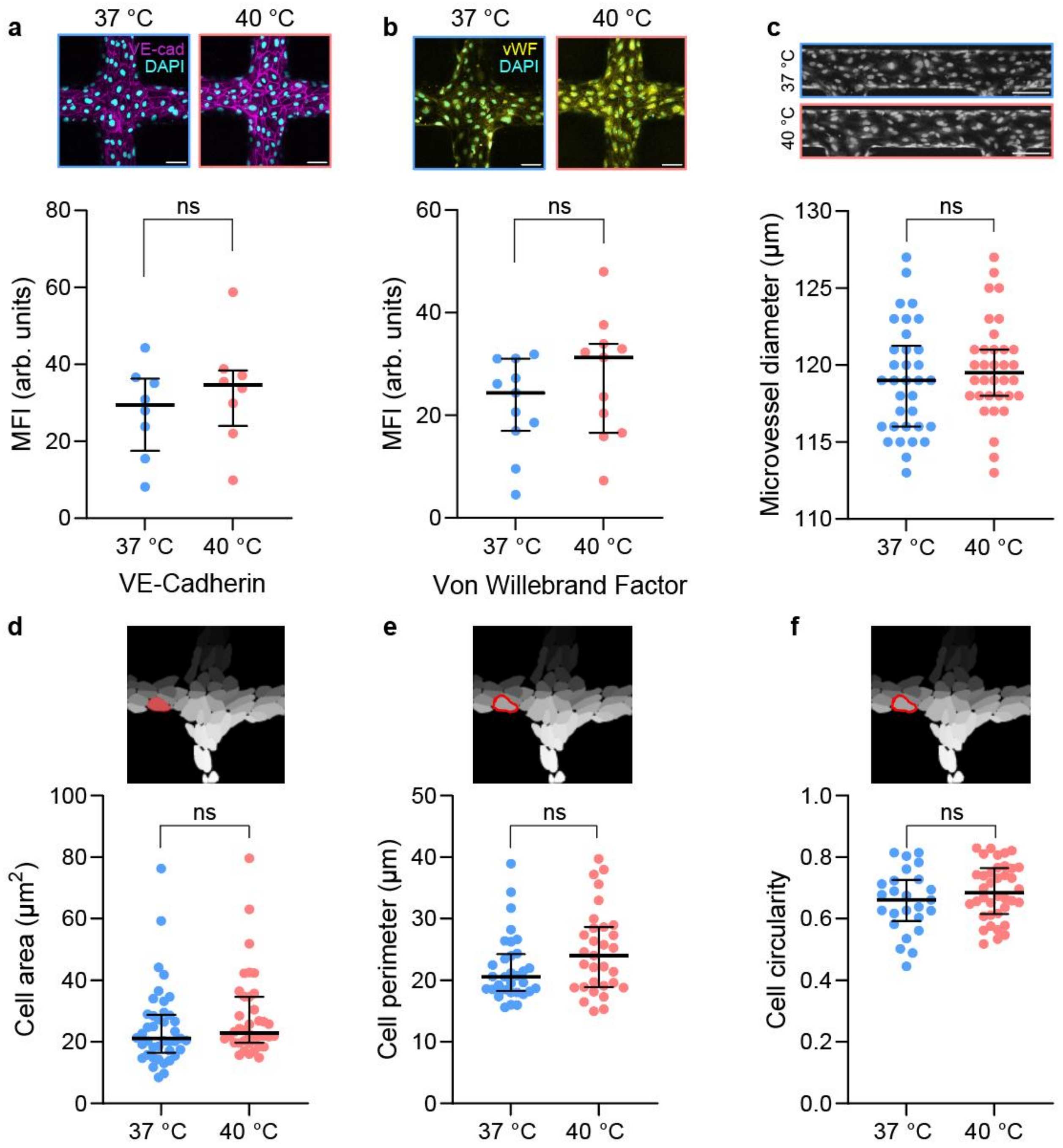


**Extended Data Figure 2. Characterisation of the effect of febrile temperature on endothelial junctions, von Willebrand Factor release and microvessel morphology.**

**a,b,** Z-projections of immunofluorescence images of 3D microvessels incubated for 1 h at either 37 °C or 40 °C and stained for **a**, VE-cadherin (magenta) or **b**, von Willebrand factor (vWF, yellow), together with DAPI (cyan) (top). Corresponding mean fluorescence intensity (MFI). *n* = 4 microvessels per condition, with each dot representing an ROI. Dot plots show mean ± s.d. Statistical analysis was performed using the Mann–Whitney *U* test. Scale bars, 50 µm.

**c,** Representative images (top) and quantification of 3D microvessel diameter (bottom) at 37 °C or 40 °C. *n* = 4 microvessels per condition, with each dot representing a microvessel edge. Dot plots show mean ± s.d. Statistical analysis was performed using the Mann–Whitney *U* test. Scale bar = 100 µm.

**d-f,** Quantification of HBMEC **d**, cell area, **e,** perimeter, and **f,** circularity in microvessels exposed to 37 °C or 40 °C. $n = 4$ microvessels per condition, with measurements taken from both the top and bottom surfaces of each microvessel. Dot plots show mean ± s.d. Statistical analysis was performed using the Mann–Whitney *U* test.

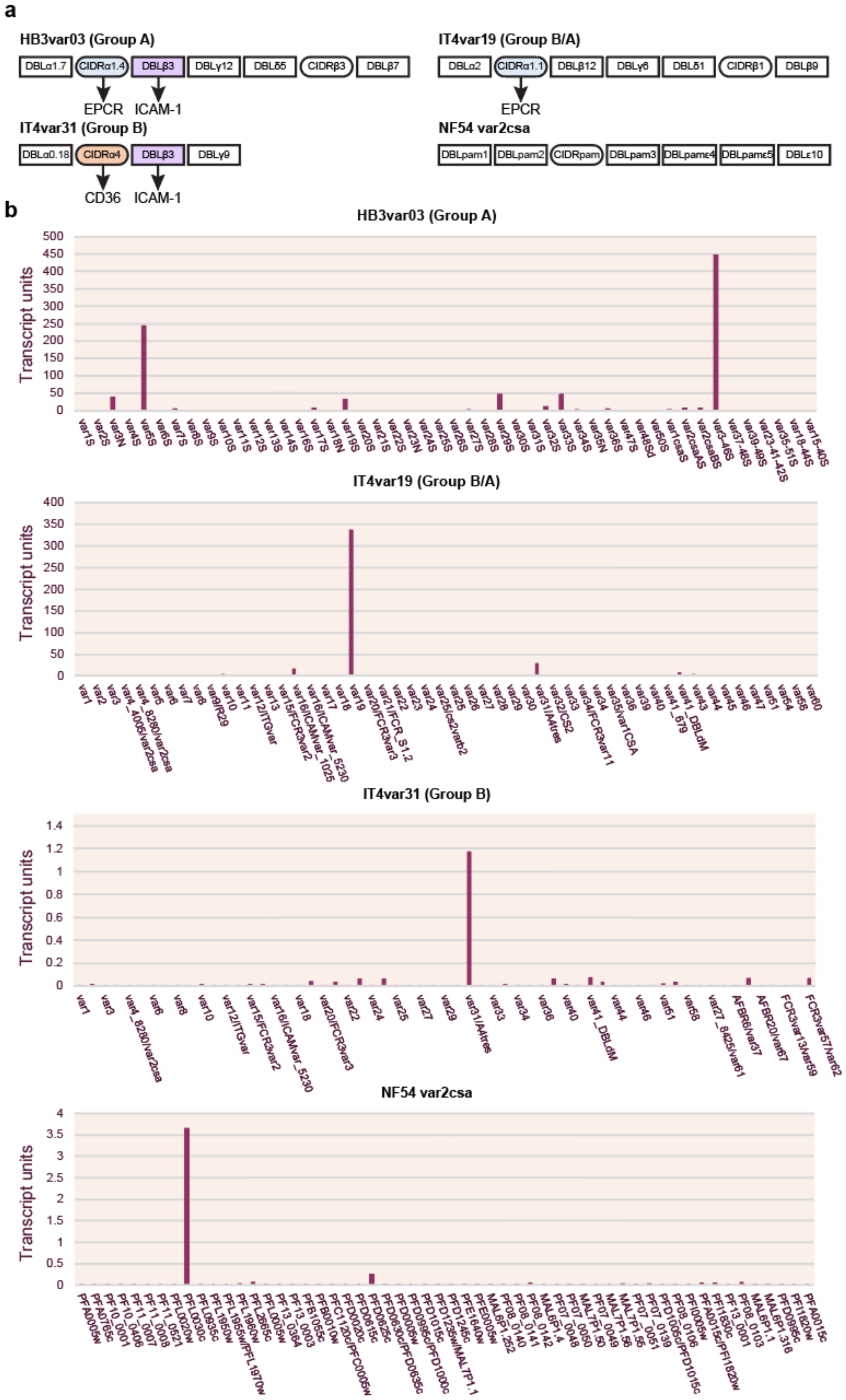
a
HB3var03 (Group A)
DBLα1.7
CIDRα1.4
DBLβ3
DBLγ12
DBLδ5
CIDRβ3
DBLβ7
EPCR
ICAM-1
IT4var19 (Group B/A)
DBLα2
CIDRα1.1
DBLβ12
DBLγ6
DBLδ1
CIDRβ1
DBLβ9
EPCR
IT4var31 (Group B)
DBLα0.18
CIDRα4
DBLβ3
DBLγ9
CD36
ICAM-1
NF54 var2csa
DBLpam1
DBLpam2
CIDRpam
DBLpam3
DBLpamε4
DBLpamε5
DBLε10
b
HB3var03 (Group A)
Transcript units
IT4var19 (Group B/A)
Transcript units
IT4var31 (Group B)
Transcript units
NF54 var2csa
Transcript units

**Extended Data Figure 3. *var* architecture and transcription profile of *Plasmodium falciparum* lines.**
**a,** Schematic representation of the PfEMP1 preferentially expressed by each experimental *P. falciparum* parasite line. Domain binding activity to endothelial receptors is indicated by arrows.
**b,** The var gene transcription profile of ring-stage iRBCs was analysed by qRT-PCR with IT4[1], HB3[2], and 3D7[3] (NF54 strain is identical to the NF54-derived clone 3D7) *var* strain-specific primer sets, as previously published[4,5]. Transcription unit levels are normalised to the housekeeping control gene STS (seryl-tRNA synthetase).

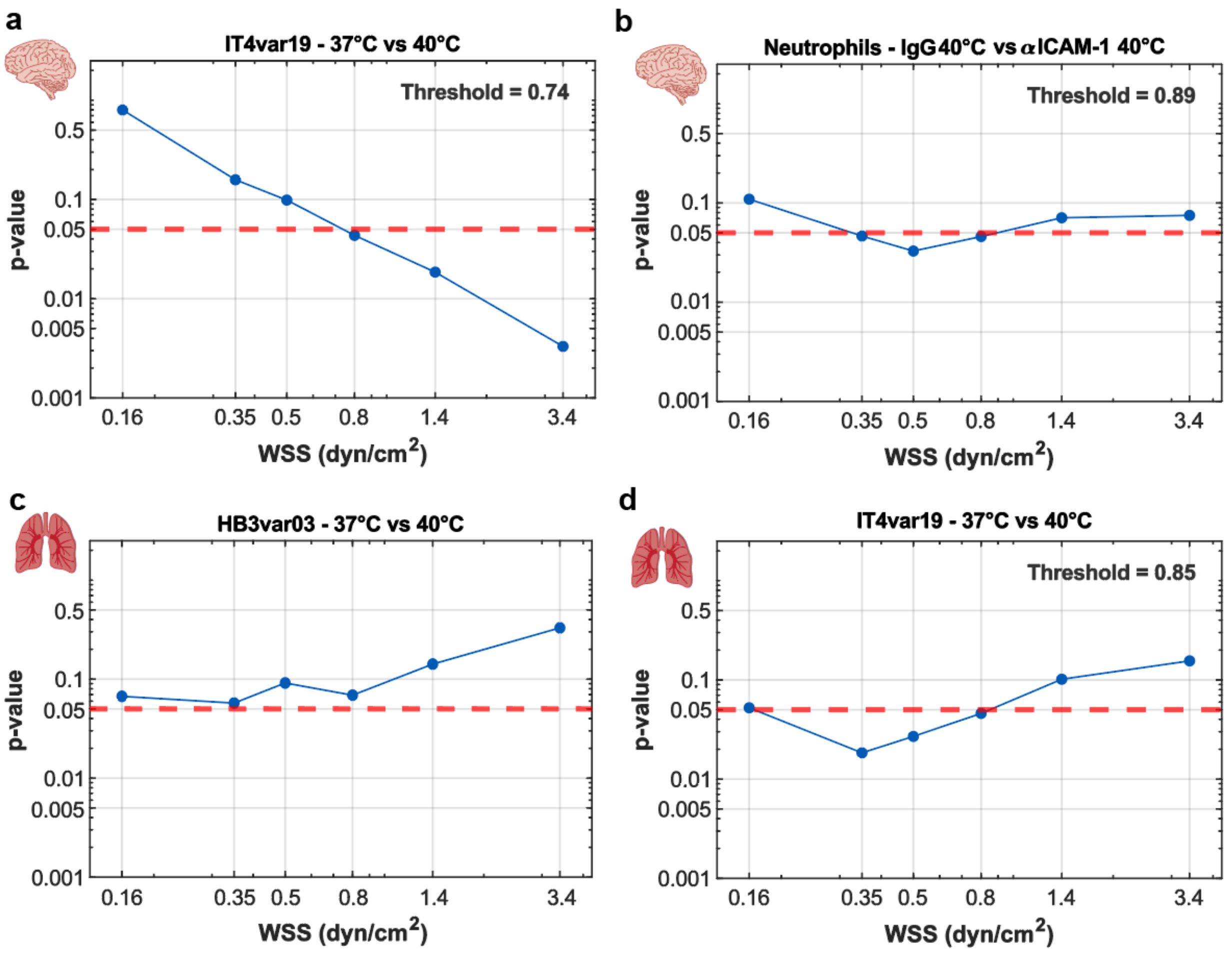


**Extended Data Figure 4 . WSS threshold sensitivity analysis.**

**a–d**, The wall shear stress (WSS) threshold was systematically varied across the observed range (0.16, 0.35, 0.5, 0.8, 1.4 and 3.4 dyn/cm$^2$), and the linear mixed-effects model (LMM) was refitted at each threshold (see Methods). Blue symbols indicate the adjusted p value obtained at each threshold, and the red dashed line indicates the significance threshold ($p = 0.05$). The threshold corresponding to the transition between statistically significant and non-significant comparisons is indicated in each panel.

**a,** IT4var19, 37 °C versus 40 °C, corresponding to Fig. 1g (3D brain microvessels).

**b,** Neutrophils, IgG versus αICAM-1 at 40 °C, corresponding to Fig. 2d (3D brain microvessels).

**c,** HB3var03, 37 °C versus 40 °C, corresponding to Fig. 4f (3D pulmonary microvessels).

**d,** IT4var19, 37 °C versus 40 °C, corresponding to Fig. 4g (3D pulmonary microvessels).

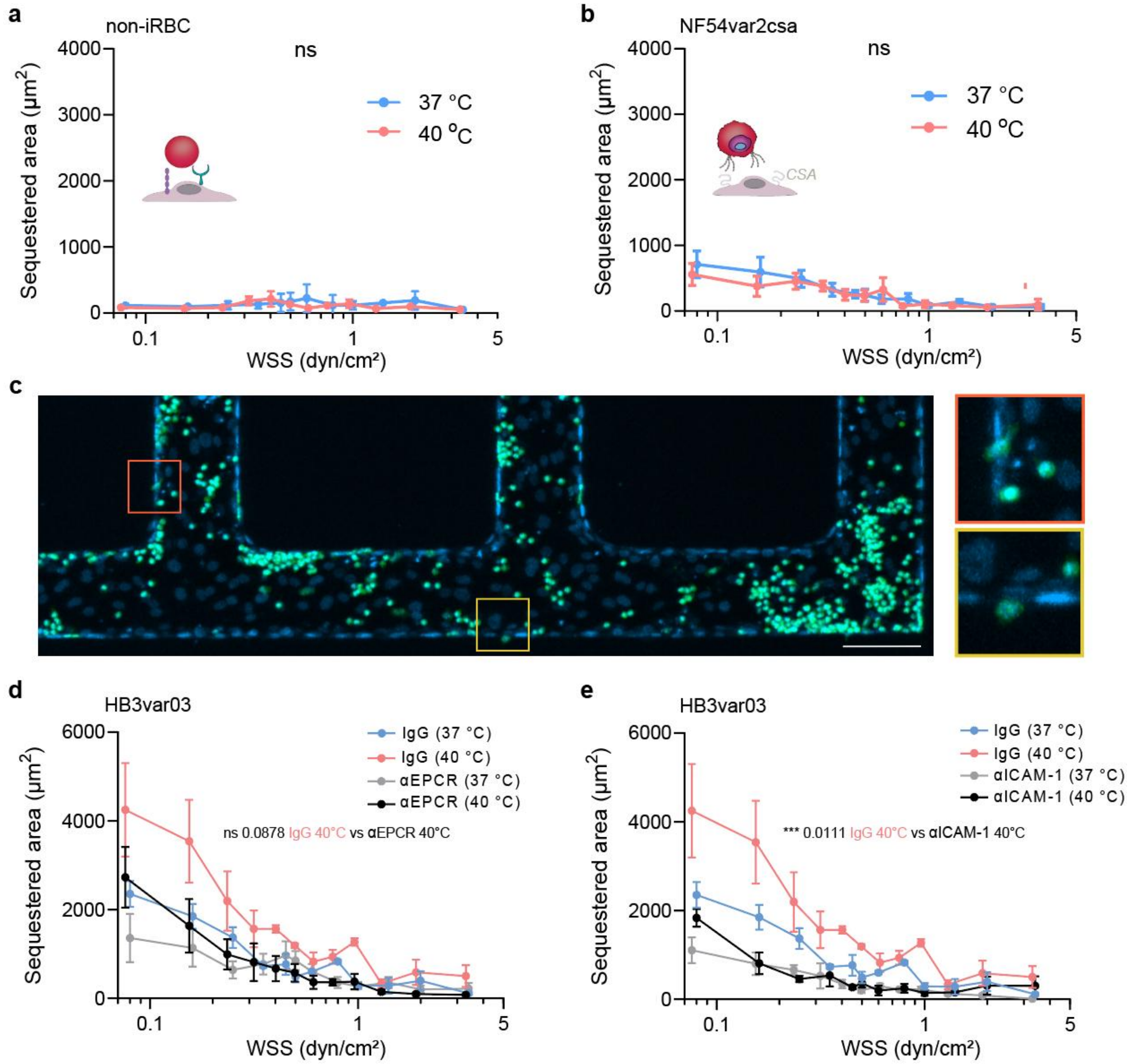


**Extended Data Figure 5. Characterisation of temperature-dependent cytoadhesion and endothelial receptor contributions**.

**a,b,** Sequestration area of non-infected RBCs (non-iRBCs; **a**) and NF54var2csa iRBCs (**b**) across the WSS range following incubation of HBMEC microvessels at 37 °C or 40 °C for 1 h. Data are shown as mean ± s.e.m.; *n* = 4 independent biological replicates. Statistical comparisons were performed using LMM accounting for WSS dependence.

**c,** Representative immunofluorescence image of a 3D microvessel showing neutrophils bound within the vessel at 40 °C. Insets highlight the occasional neutrophils appearing to initiate transmigration (red box) or located outside the vessel (yellow box), whereas the majority of neutrophils remained within the vessel lumen. Scale bar, 100 µm.

**d,** HB3var03 iRBC sequestration area across the WSS range following 1 h incubation of microvessels at 37 °C or 40 °C in the presence of IgG isotype control or anti-EPCR mAb 252.

**e,** HB3var03 iRBC sequestration area across the WSS range following 1 h incubation of microvessels at 37 °C or 40 °C in the presence of IgG isotype control or anti-ICAM-1 mAb 15.2. Data in **d,e** are mean ± s.e.m.; *n* = 4–6 independent biological replicates. Statistical comparisons were performed using LMM accounting for WSS dependence.

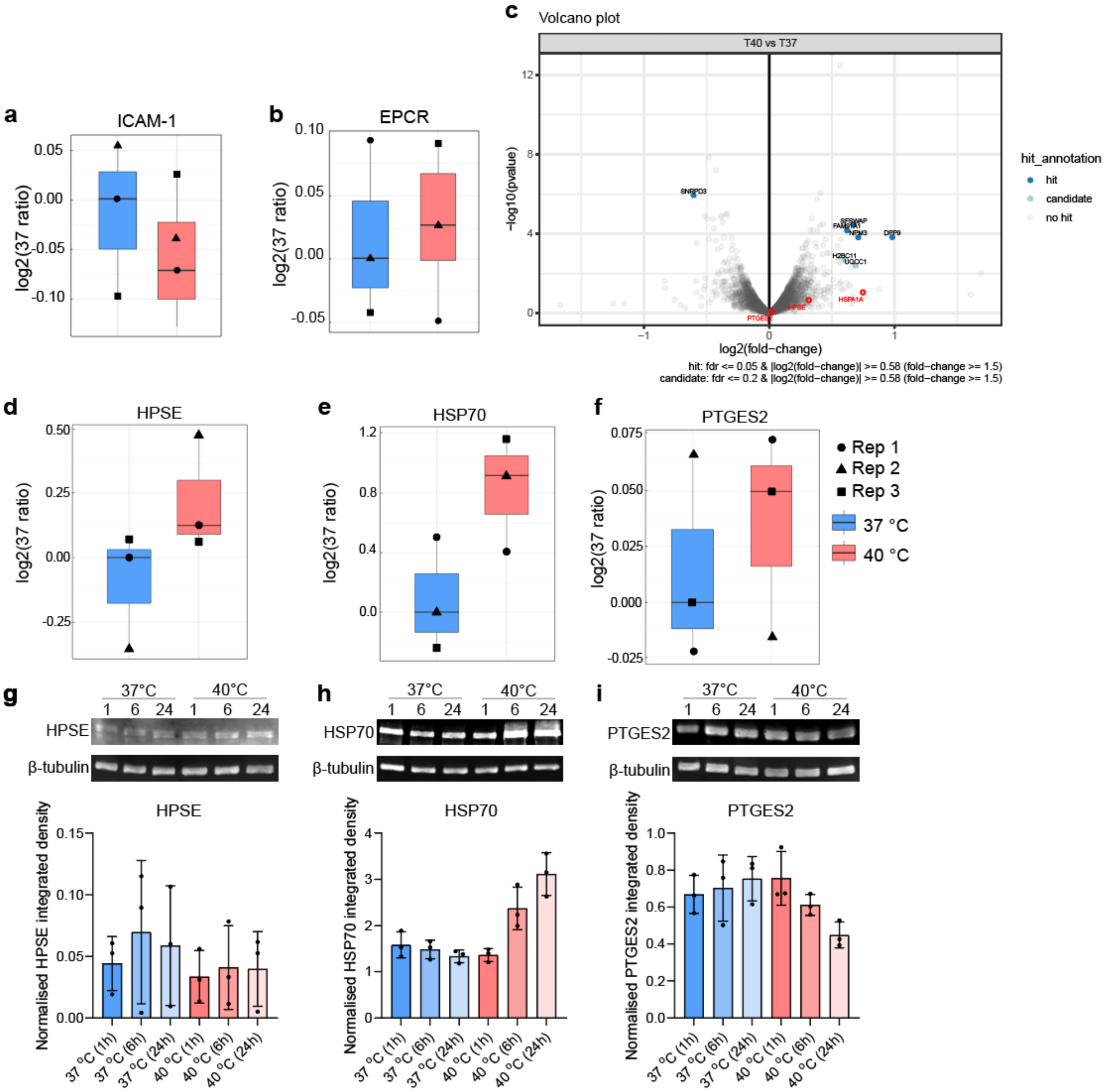


**Extended Data Figure 6. Proteomic analysis and validation of endothelial proteins associated with febrile temperature.**

**a–f,** Proteomic analysis of HBMECs exposed to 37 °C or 40 °C for 1 h.

**a,b,** Relative abundance of **a**, ICAM-1 and **b**, EPCR.

**c,** Volcano plot showing differential protein abundance between HBMECs exposed to 40 °C and 37 °C. Proteins meeting the predefined criteria for differential abundance are highlighted. Heparanase (HPSE), heat shock protein 70 (HSPA1A) and prostaglandin E synthase 2 (PTGES2) are additionally highlighted because of their relevance to glycocalyx shedding and the heat-shock response.

**d–f**, Relative abundance of **d**, HPSE, **e**, HSPA1A, and **f**, PTGES2. Protein abundance in **a,b, d,e,f** is shown as the log2-transformed ratio of normalised TMT reporter ion intensity relative to the median value at 37 °C. Each point represents one independent biological replicate ($n$ = 3).

**g–i**, Western blot validation of **g**, HPSE, **h**, HSP70, and **i**, PTGS2 protein abundance following exposure to 37 °C or 40 °C for 1, 6 or 24 h. $n$ = 3 independent biological replicates. Data are shown as mean ± s.e.m.; Representative western blot images are shown alongside the corresponding quantification. Statistical comparisons were performed using two-sided Welch's $t$-tests.

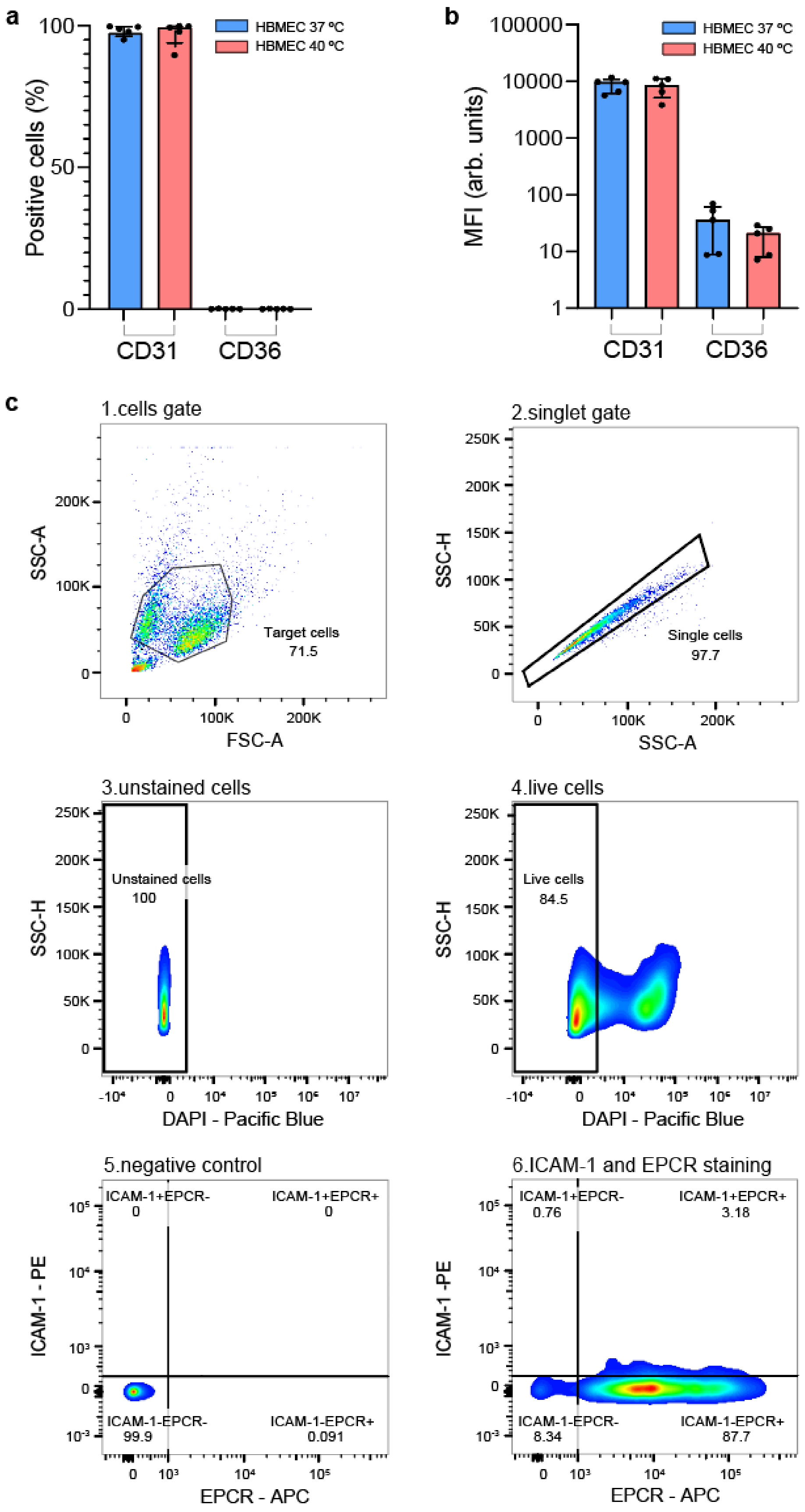
a
Positive cells (%)
100
50
0
HBMEC 37 ºC
HBMEC 40 ºC
CD31
CD36
b
MFI (arb. units)
100000
10000
1000
100
10
1
HBMEC 37 ºC
HBMEC 40 ºC
CD31
CD36
c
1.cells gate
SSC-A
FSC-A
Target cells
71.5
2.singlet gate
SSC-H
SSC-A
Single cells
97.7
3.unstained cells
SSC-H
DAPI - Pacific Blue
Unstained cells
100
4.live cells
SSC-H
DAPI - Pacific Blue
Live cells
84.5
5.negative control
ICAM-1 - PE
EPCR - APC
ICAM-1+EPCR-
0
ICAM-1+EPCR+
0
ICAM-1-EPCR-
99.9
ICAM-1-EPCR+
0.091
6.ICAM-1 and EPCR staining
ICAM-1 -PE
EPCR - APC
ICAM-1+EPCR-
0.76
ICAM-1+EPCR+
3.18
ICAM-1-EPCR-
8.34
ICAM-1-EPCR+
87.7

**Extended Data Figure 7. HBMEC surface expression of endothelial receptors.**
**a,b,** Percentage of HBMECs expressing surface CD31 and CD36 and corresponding mean fluorescence intensity (MFI) following 1 h incubation at 37 °C or 40 °C. $n = 5$ independent biological replicates Bars represent mean ± s.d.. Statistical comparisons were performed using the Mann–Whitney *U* test.
**c,** Representative flow cytometry gating strategy for HBMECs, showing sequential gating of 1. target cells, 2. singlets, 3. unstained and 4. live cells selected by DAPI staining, followed by 5. negative control and 6. double ICAM-1 and EPCR stained cells (see details in Methods).

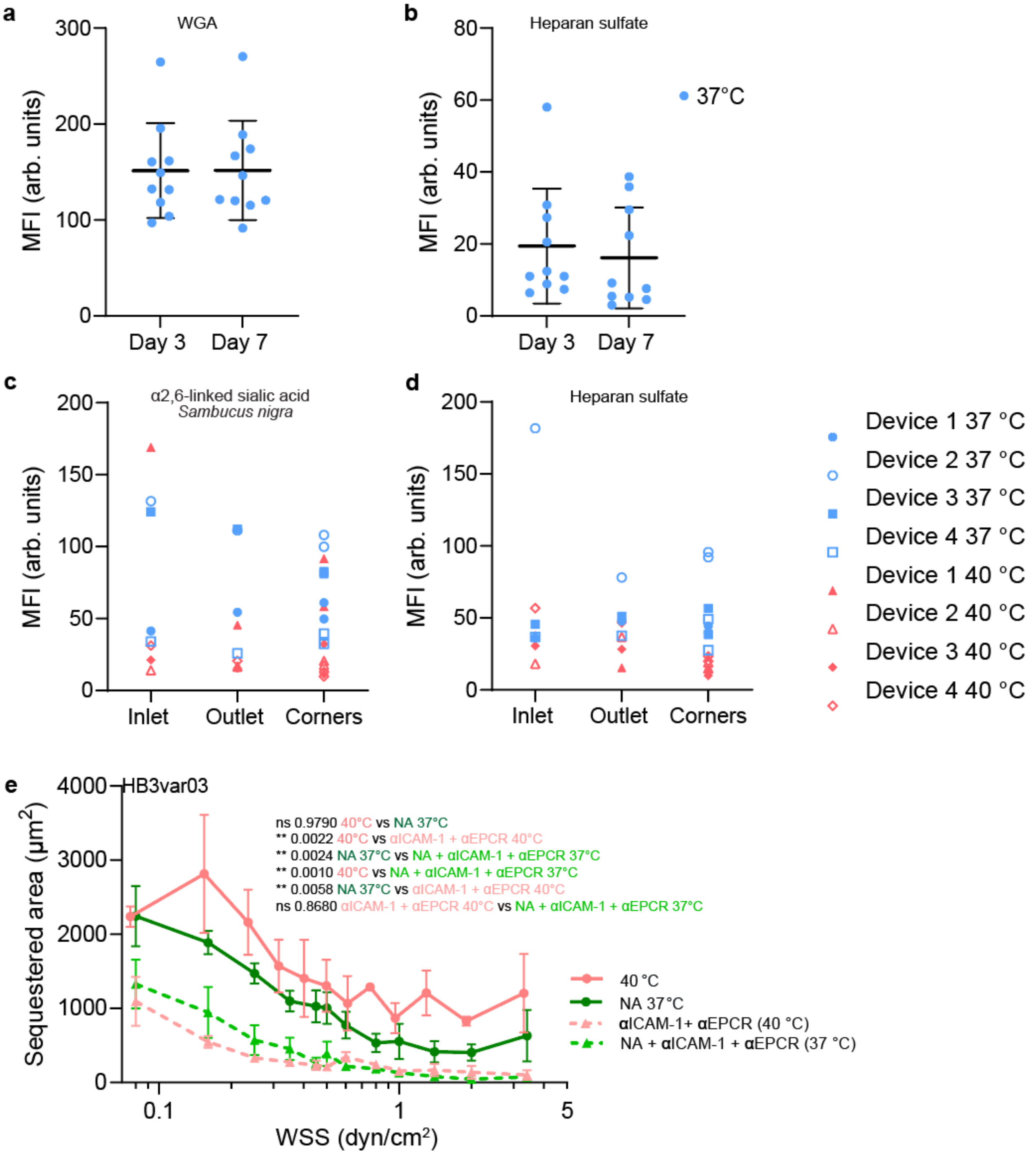


**Extended Data Figure 8. Glycocalyx maturation and contribution of EPCR and ICAM-1 to iRBC adhesion following sialic acid disruption by neuraminidase.**

**a,b,** Mean fluorescence intensity (MFI) of FITC-conjugated wheat germ agglutinin (WGA) (**a**) and heparan sulfate (**b**) staining in 3D microvessels after 3 or 7 days of gravity-driven perfusion. $n = 3$ microvessels per condition, with each dot representing an ROI (mean ± s.d.). Statistical analysis was done using the Mann–Whitney *U* test.

**c,d,** MFI of α2,6-linked sialic acid stained with *Sambucus nigra* lectin (**c**) and heparan sulfate (**d**) measured in inlet, outlet and corner regions of microvascular networks following 3 days of gravity-driven perfusion and subsequent 1 h incubation at 37 °C or 40 °C. $n = 4$ independent biological replicates per condition.

**e,** HB3var03 iRBC sequestration area across the WSS range following incubation of microvessels at 40 °C or treatment with neuraminidase (NA) at 37 °C, in the presence or absence of anti-ICAM-1 mAb 15.2 and anti-EPCR mAb 252. *n* = 4–6 independent biological replicates (mean ± s.e.m.). Statistical comparisons in e were performed using LMM accounting for WSS dependence, with pairwise comparisons adjusted for multiple testing.

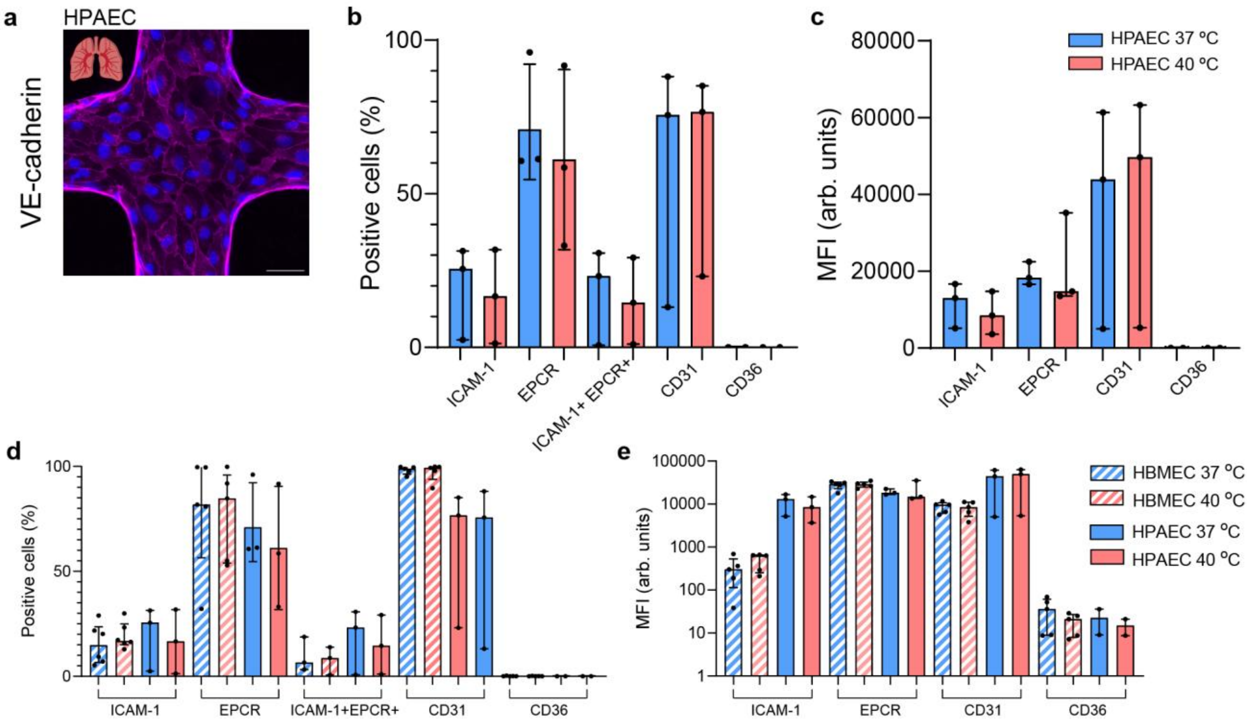


**Extended Data Figure 9. Endothelial VE-cadherin and receptor expression in pulmonary microvessels at 37 °C and 40 °C.**

**a,** Representative z-stack projection immunofluorescence image of a cross-section through a bioengineered pulmonary microvessel (human pulmonary artery endothelial cells, HPAEC), stained for VE-cadherin (magenta) and nuclei (DAPI, blue). Scale bar, 50 µm.

**b,c,** Percentage of HPAECs positive for surface ICAM-1+, EPCR+, ICAM-1$^{+}$EPCR$^{+}$ double-positive cells, CD31+, and CD36+, and corresponding mean fluorescence intensity (MFI) for ICAM-1, EPCR, CD31, and CD36, following 1 h incubation at 37 °C or 40 °C. $n$ = 3 independent biological replicates. Bars represent mean ± s.d.; pairwise statistical comparisons were performed using the Mann–Whitney *U* test.

**d,e,** Comparison of the percentage of positive cells (**d**) and MFI (**e**) between HBMECs and HPAECs following 1 h incubation at 37 °C or 40 °C. $n$ = 3 for HPAECs and $n$ = 5–6 for HBMECs. Bars represent mean ± s.d. Pairwise statistical comparisons were performed using the Mann–Whitney *U* test.

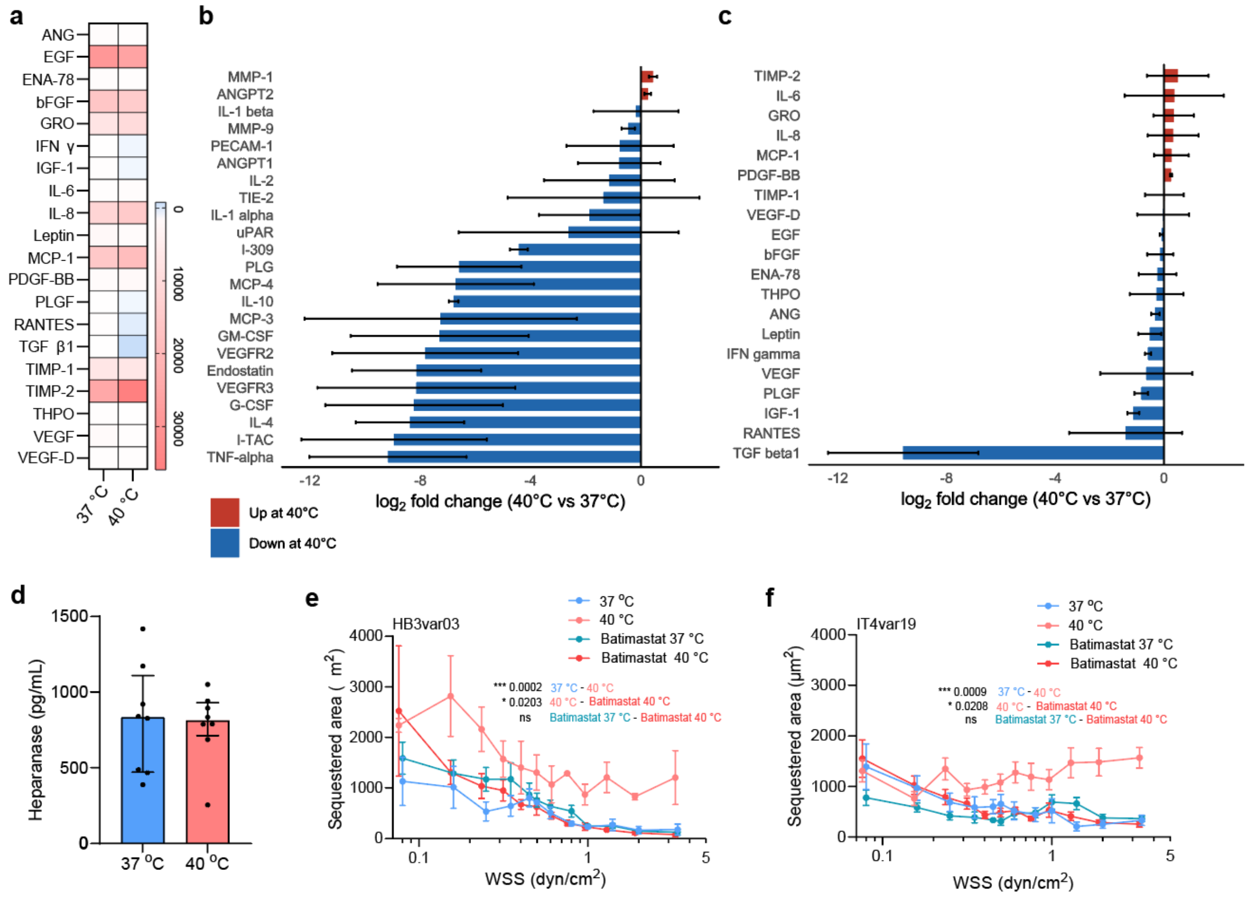


**Extended Data Figure 10. Secreted mediators of glycocalyx degradation and effect of MMP inhibition on cytoadhesion induced by febrile-temperature.**

**a**, Heatmap showing semiquantitative protein array signal intensities of angiogenic mediators, inflammatory cytokines, and proteolytic enzymes secreted by HBMEC monolayers following 1 h incubation at 37 °C or 40 °C.

**b,c**, Corresponding $log_2$ fold change (37 °C vs 40 °C) for the same analytes, shown across two array panels (a, and Fig. 5a in the manuscript); red, increased at 40 °C; blue, decreased at 40 °C. MMP-1 $p$ value = 0.031 was assessed using the paired Wilcoxon signed-rank; Benjamini–Hochberg (BH)-adjusted $P$ = 0.18 and ANGPT2, $p$ = 0.1563; BH-adjusted $p$ = 0.4799 across the 23-protein panel. TIMP-2, $p$ = 0.3125; BH-adjusted $p$ = 0.568 across the 20-protein panel. $n$ = 3 independent biological replicates. Bars represent mean log2 fold change (40 °C vs 37 °C) ± s.d..

**d**, Heparanase concentration in HBMEC supernatants following 1 h incubation at 37 °C or 40 °C, measured by ELISA. $n$ = 8 independent biological replicates. Bars represent median and interquartile range. Statistical comparisons were performed using two-sided Welch's $t$-tests.

**e,f**, Sequestered area of HB3var03- (**e**) and IT4var19- (**f**) iRBCs across the WSS range, at 37 °C, 40 °C, or following pre-treatment with the broad-spectrum MMP inhibitor batimastat at 37 °C or 40 °C. $n$ = 6 independent biological replicates (mean ± s.e.m.). Statistical comparisons were performed using LMM accounting for WSS dependence, with pairwise comparisons adjusted for multiple testing.
